# Comparison of Automated White Matter Lesion Segmentation Approaches for Use in Large, Multi-Site Data Analyses in Parkinson’s Disease

**DOI:** 10.64898/2026.05.27.726795

**Authors:** Sarah Al-Bachari, So Hoon Yoon, Phoebe Emson, Shauna Angell, John Cain, Aswin Abraham, Azeem Chughtai, Edward Sizer, Edwin Barnes, Maryam Al-Wardy, Siddarth Kannan, Rohan Paul-Thaper, Joanna Bright, Conor Owens-Walton, Corey T. McMillan, Johannes C. Klein, Ludovica Griffanti, Sophia I. Thomopoulos, Neda Jahanshad, Paul M. Thompson, Ysbrand D. van der Werf, Chris Vriend, Laura M. Parkes, Hedley C.A. Emsley, Anette Schrag, Hamied A. Haroon

## Abstract

**Background:** Parkinson’s disease (PD) is the second most common neurodegenerative disorder. PD currently lacks effective disease-modifying treatments, likely due to its diverse clinical features and underlying neuropathology. The vascular role in PD is emerging, with vascular mechanisms increasingly implicated, yet the literature remains conflicted, motivating large-data analyses with greater statistical power. White matter lesions (WML) are an accepted imaging marker of small vessel disease. Accurate automated WML segmentation techniques are crucial for large-scale studies in PD due to the impracticality of manual segmentation for extensive datasets and to ensure consistency. Evaluation of the optimum approach in PD for large-scale analysis is lacking. This study aimed to evaluate various automated WML segmentation algorithms to determine the most accurate and reliable method, among those selected, for assessing WML for multi-site large data analysis in PD.

**Methods:** We assessed whole-brain volumetric T1-weighted and FLAIR images from 201 PD patients (mean age, 66.6 ± 7.86 years) and 64 healthy controls (HC; mean age, 66.3 ± 8.67) across three datasets: the Parkinson’s Progression Markers Initiative (PPMI), the University of Pennsylvania (UPenn) and the Montreal Neurological Institute Biobank: Clinical Biological Imaging and Genetic Repository (C-BIG). The sample included different scanners, imaging parameters and lesion loads, as would be expected for multi-site data. WML were manually segmented to provide the gold standard, and four freely available automated algorithms were evaluated: FSL’s BIANCA, FreeSurfer, SPM’s LST-LPA and U-Net-pgs using the performance metrics: Dice score, Hausdorff distance, recall, precision, F1 score, log absolute volume difference (LOGAVD) and intraclass correlation coefficient (ICC). Subgroup analyses were performed based on lesion load and lobar regions. The associations of data from these automated approaches with age, and with Fazekas and Wahlund visual rating scales, were assessed through partial correlation analysis.

**Results:** U-Net-pgs performed best overall, with the highest Dice score (PD: 0.46 ± 0.21; HC: 0.39 ± 0.21), recall (PD: 0.76 ± 0.25; HC: 0.62 ± 0.31), precision (PD: 0.49 ± 0.25; HC: 0.63 ± 0.27), F1 score (PD: 0.54 ± 0.22; HC: 0.56 ± 0.22) and ICC (PD: 0.965; HC: 0.967) and lowest Hausdorff distance (PD: 8.89 ± 3.96; HC: 6.33 ± 2.91). U-Net-pgs achieved the lowest LOGAVD in the PD group (0.31 ± 0.31) whereas BIANCA-LOO with a threshold of 0.9 was lowest in HC (0.27 ± 0.30). U-Net also showed superior performances in all lesion loads for PD and overall across various brain regions in both PD and HC.

**Conclusion:** Overall, U-Net-pgs emerged as the best performing automated method, of those we evaluated, for WML segmentation in PD and HC within a dataset collected with various scanner and image acquisition parameters. U-Net-pgs consistently outperformed other automated approaches across lesion loads and brain regions, for most metrics. The accuracy and reliability of U-Net-pgs make it a promising tool for large-scale analyses, facilitating future research investigating WML in PD.

## INTRODUCTION

Parkinson’s disease (PD) is now recognized as the fastest-growing neurological disorder worldwide in terms of prevalence, disability and societal burden. About 6.1 million individuals globally had PD in 2016, and this number is anticipated to surpass 14 million by the year 2040 (Dorsey et al., 2018). PD is not only highly variable in terms of clinical presentation, with over 50 possible symptoms, but also in underlying neurobiology (Espay et al., 2017; Mestre et al., 2021). Within this diversity, there is an urgent need to identify groups who will benefit from specific treatments, i.e., a move towards a precision medicine approach (Thenganatt and Jankovic, 2014; Greenland et al., 2019; Marras et al., 2020; Hu et al., 2024). Due to the heterogeneity in PD, large-scale data analysis is required to enable subtyping and identify at-risk groups who may be responsive to certain treatments.

The vascular role in PD is rapidly emerging. The biological underpinnings lie at the level of the neurovascular unit, with blood-brain barrier dysfunction and cerebral hypoperfusion interacting with the other key pathogenic processes that contribute to selective neuronal loss (Jellinger, 2003; Zlokovic, 2008; Al-Bachari et al., 2017; Sweeney et al., 2018; Al-Bachari et al., 2020). Large, multi-center longitudinal studies of PD have demonstrated cerebral vascular changes associated with greater PD-related disability and more severe gait, cognitive and mood phenotypes, associated with greater disability that tend to be less responsive to levodopa treatment (Stojkovic et al., 2018; Chen et al., 2020; Lee et al., 2020). These changes may not only accompany neurodegeneration but actively contribute to it, holding promise as a target for a disease-modifying treatment (Monti et al., 2022; Paul and Elabi, 2022; Kieliszek-Ryba et al., 2025), yet a vascular biomarker is lacking.

MRI markers of small vessel disease include white matter lesions (WML), lacunes, enlarged perivascular spaces, and cerebral microbleeds (Wardlaw et al., 2013; Chojdak-Łukasiewicz et al., 2021). Of these, WML are the most commonly measured marker and are particularly relevant in PD as they are associated with motor and cognitive deficits (Joutel and Chabriat, 2017; Duering et al., 2023). WML appear as hyperintensities on T2-weighted fluid-attenuated inversion recovery (FLAIR), while they appear as hypointensities on T1-weighted (T1w) MRI. Their measurement is non-invasive and highly reproducible, making WML an ideal neuroimaging marker for studies in PD. Several studies have demonstrated that higher WML burden correlates with worse outcomes in PD, including cognitive decline, gait disturbances, and increased risk of dementia (González-Redondo et al., 2012; Kandiah et al., 2013; Sunwoo et al., 2014; Compta et al., 2016; Malek et al., 2016; Ciliz et al., 2018; Stojkovic et al., 2018; Pozorski et al., 2019; Wu et al., 2023; Zou et al., 2023). However, despite over a decade of research, there are conflicting results, with some studies showing no association of WML with clinical features or disease severity in PD (Dalaker et al., 2009a, 2009b; Despotović et al., 2015; Lee and Lee, 2016; Hanning et al., 2019). This discrepancy is likely due to both the heterogeneity of PD pathophysiology and methodological differences in identifying WML. These include small sample sizes, differences in analysis methods, and varying sensitivity of approaches.

Manual methods remain the current gold standard for WML segmentation; however, they are labor intensive, time consuming and show inter-rater variability. Thus, it is imperative to identify an automated algorithm for quantifying WML burden and location accurately for use in large-scale analyses (Hotz et al., 2022). Numerous automated WML segmentation algorithms have been proposed, including supervised, unsupervised, and semi-supervised methods depending on the quantity of expertly annotated data available. These methods have been extensively described elsewhere, yet in summary each technique has its advantages and disadvantages, including poor performance in low lesion loads, requiring a high degree of human intervention and expertise, the requirement of a training set and varying false positive and false negative rates (Caligiuri et al., 2015). Few studies have compared the performance and validity of these different methods.

MRI studies are often small and fail to account for the heterogeneity of PD. Pooling data from multiple cohorts, often using consortia data, aims to overcome this challenge (Thompson et al., 2020). Consortium data have contributed significantly to our understanding of various disease processes (Weaver et al., 2021; Chen et al., 2025), yet certain challenges still prevail, namely differences in study protocols including scanner parameters, imaging acquisitions and clinical data collected (Zugman et al., 2022; Giehl et al., 2024). Therefore, imaging analysis methods that can withstand variations in scanner and image acquisition parameters need to be sought to allow for more uniform analysis in pooled datasets.

We here compared several freely available automated WML segmentation algorithms that have recently shown promise in producing accurate WML segmentations mostly in healthy aging and cognitively impaired populations (Heinen et al., 2019; Kuijf et al., 2019; Vanderbecq et al., 2020; Hotz et al., 2022; Gaubert et al., 2023; Torres-Simon et al., 2024). These included: FSL’s Brain Intensity AbNormality Classification Algorithm (BIANCA) (Griffanti et al., 2016), FreeSurfer (Fischl, 2012), SPM’s Lesion Segmentation Tool Lesion Prediction Algorithm (LST-LPA) (Schmidt et al., 2012) and UNet-pgs (Ronneberger et al., 2015; Li et al., 2018; Park et al., 2021). These approaches are summarized in **Table 1** below. To the authors’ knowledge, these approaches have not been tested specifically to assess WML in PD, nor in PD consortium data that includes different scanners and variable imaging parameters such as slice thickness, voxel size and field strength, all of which may impact the accuracy of WML segmentations (Khademi et al., 2021; Tran, 2022).

**Table 1:** Summary of the automated approaches for WML segmentation used in this study.

| Name | Summary |
| --- | --- |
| **BIANCA**<br>(Griffanti et al., 2016) | BIANCA is a supervised WML segmentation tool that applies a k-nearest neighbor classifier. It requires a training set of manually segmented lesion masks which are used to determine the similarity between features of the target voxel and those in the training set to estimate the lesion probability for each voxel. This results in a probabilistic map of WML; a threshold can be applied to binarize these. |
| **FreeSurfer**<br>(http://surfer.nmr.mgh.harvard.edu/) | FreeSurfer applies an automated pipeline to the T1w image including intensity normalization, brain extraction, surface reconstruction and volumetric labelling steps to delineate cortical and subcortical structures. A probabilistic anatomical atlas derived from manually annotated MRI datasets is used to assign each voxel to the most likely tissue class based on spatial context and intensity features. Results include a binary map of white matter hypointensities from the T1 image. |
| **LST-LPA (LPA)**<br>(www.statistical-modelling.de/lst.html; Schmidt et al., 2012) | LST-LPA uses a logistic-regression classifier trained on FLAIR images. A lesion belief map is generated from the training set and used to estimate the lesion probability at each voxel. The spatial information is combined with image intensity information to form a lesion map for each subject. This results in a probabilistic map of WML and a threshold can be applied to binarize these. |
| **UNet-pgs (U-Net)**<br>(Li et al., 2018; Park et al., 2021) | UNet-pgs (referred to simply as U-Net in this paper) is a supervised convolutional neural network approach which can be applied to medical image segmentation. It is trained on co-registered T1w and FLAIR images using highlighting foregrounds to improve sensitivity for small or boundary lesions affected by partial volume effects. This results in a binary map of WML. |

The aim of this study was to compare and identify an optimal automated WML segmentation approach for large multi-site data in PD. The following questions were to be answered by this study:

1. Assessment of segmentation performance for use in large-scale multi-site data: *Which automated approach provides estimates that are most consistent with the gold standard of manual segmentations, assessed by established performance metrics* (Kuijf et al., 2019)?
2. Validations using visual rating scales and age: *How do the WML segmentations provided by the different approaches relate to Fazekas and Wahlund scores and chronological age?*

## MATERIALS AND METHODS

### Participants

Demographic and brain MRI data were gathered from three datasets: the Montreal Neurological Institute Biobank: Clinical Biological Imaging and Genetic Repository (C-BIG, https://pmc.ncbi.nlm.nih.gov/articles/PMC9537233/) (Das et al., 2022), the Parkinson’s Progression Markers Initiative (PPMI, https://www.ppmi-info.org/) and the University of Pennsylvania (UPenn, https://www.med.upenn.edu/pdmdc/). All participants underwent comprehensive clinical and imaging assessments. Written informed consent for participation and sharing of anonymized data was obtained from all participants at the relevant site. PD diagnosis was based on the UK Parkinson’s Disease Brain Bank criteria. Participants with any other significant CNS disorders were excluded. Healthy controls (HC) did not have PD or significant other CNS disorders.

### MRI Data Acquisition

All participants underwent 1.5T or 3T 2D or 3D FLAIR and 3D T1w imaging. Scanner parameters including voxel sizes, repetition times, echo times and slice thickness varied [**Table 4**]. All images were visually inspected for quality to confirm that no motion artifacts or poor image quality could substantially impact segmentations.

### Manual Segmentation of WML

WML were manually segmented to establish a gold standard for assessing the performance of WML segmentation by automated methods. FLAIR images for each participant were manually segmented by two trained raters (PE and SA) in the axial plane using ITK-SNAP software. PE and SA were both trained by consultant neuroradiologist JC. All manual segmentations were performed blinded to the automated segmentations.

### Automated WML Segmentation Algorithms

The following automated approaches were applied to the complete dataset: BIANCA, FreeSurfer, LPA and U-Net. The specific efforts made to optimize the approaches are outlined below.

#### BIANCA

Both T1w and FLAIR images were used as inputs to maximize classifier performance. Bias field correction was applied to the images and the T1w image registered to the FLAIR image using the FSL tools FMRIB’s Automated Segmentation Tool (FAST) and the Linear Image Registration Tool, respectively (Zhang et al., 2001; Jenkinson et al., 2012).

Two training strategies were applied. A leave-one-out approach (BIANCA-LOO), segmented each subject’s image using all remaining subjects as the reference set. This was done for both PD and HC separately and required all data to be manually labelled. As this approach would not be practical for unseen multi-site data, a trained-model approach (BIANCA-training), which generated training data from two of the three datasets then was applied to the third dataset, was also evaluated. Due to the low number of HC, this was only done in PD.

Binary lesion masks were generated using thresholds of 0.9 and 0.99 (BIANCA (0.9), BIANCA (0.99)), based on in-house threshold testing, visual inspection of the outputs and recommended optimal thresholds (Griffanti et al., 2016)

#### FreeSurfer

T1w images were used as the inputs for FreeSurfer (version 7.2.0). To optimize the approach, bias field correction using FMRIB’s Automated Segmentation Tool (FAST) was applied (Zhang et al., 2001). Both bias field corrected (FreeSurfer-FAST) and uncorrected (FreeSurfer) approaches were used in the study. FreeSurfer’s tissue segmentation and parcellation pipeline was used with default settings to identify white matter hypointensities from T1w images which were assumed to reflect WML.

The outputs in T1w space were transformed to FLAIR space using FSL (Jenkinson et al., 2012) to allow comparisons with the manual segmentations in FLAIR space. The co-registered masks were visually inspected to ensure successful co-registration and ensure spatial agreement between manual and automated segmentations.

#### LPA

Both T1w and FLAIR images were used as inputs to maximize classifier performance (version 3.0.0 of the LST toolbox (www.statistical-modelling.de/lst.html) for SPM12 in MATLAB (R2025a, The Mathworks)). Bias field correction and transformation of the T1w image to FLAIR space as described for FreeSurfer above was also applied. Both bias field corrected (LPA-FAST) and uncorrected (LPA) approaches were used in the study.

To generate binary lesion masks, thresholds of 0.4 and 0.5 (LPA (0.4), LPA (0.5)) were applied to the probability maps based on in-house threshold testing, visual inspection of the outputs and previous studies (Torres-Simon et al., 2024).

#### U-Net

We adapted a pre-processing step to register the FLAIR to the T1w rather than vice versa due to T1w images often demonstrating higher anatomical resolution and contrast. The outputs in T1w space were transformed to FLAIR space using FSL (Jenkinson et al., 2012) as above.

### Grouping According to WML Loads

The WML load was determined from the volume of the manual segmentations using ITK-SNAP (Yushkevich et al., 2006). All participants were divided into three groups based on the load of WML. Participants with WML loads less than 5 cm^3^ were grouped into the ‘low WML load group’, those with more than 15 cm^3^ were grouped into the ‘high WML load group’ and the rest were grouped into the ‘medium WML load group’ (Griffanti et al., 2016; De Sitter et al., 2017; Ong et al., 2022). This classification was based on the total lesion volume regardless of whether the WML were contiguous or sparse. The performance of each automated WML approach was analyzed using the performance metrics below according to these subgroups in PD and in the low WML load group in HC.

### Performance Metrics

All metrics were computed using in-house code in MATLAB (R2025a, The Mathworks) and Python (version 3.8, Python Software Foundation). The metrics used are summarized in **Table 2**. This combination of metrics was selected to not only ensure volumetric and spatial agreement between the automated approaches and manual segmentation, but to understand the influence of true/false positives/negatives.

**Table 2:**
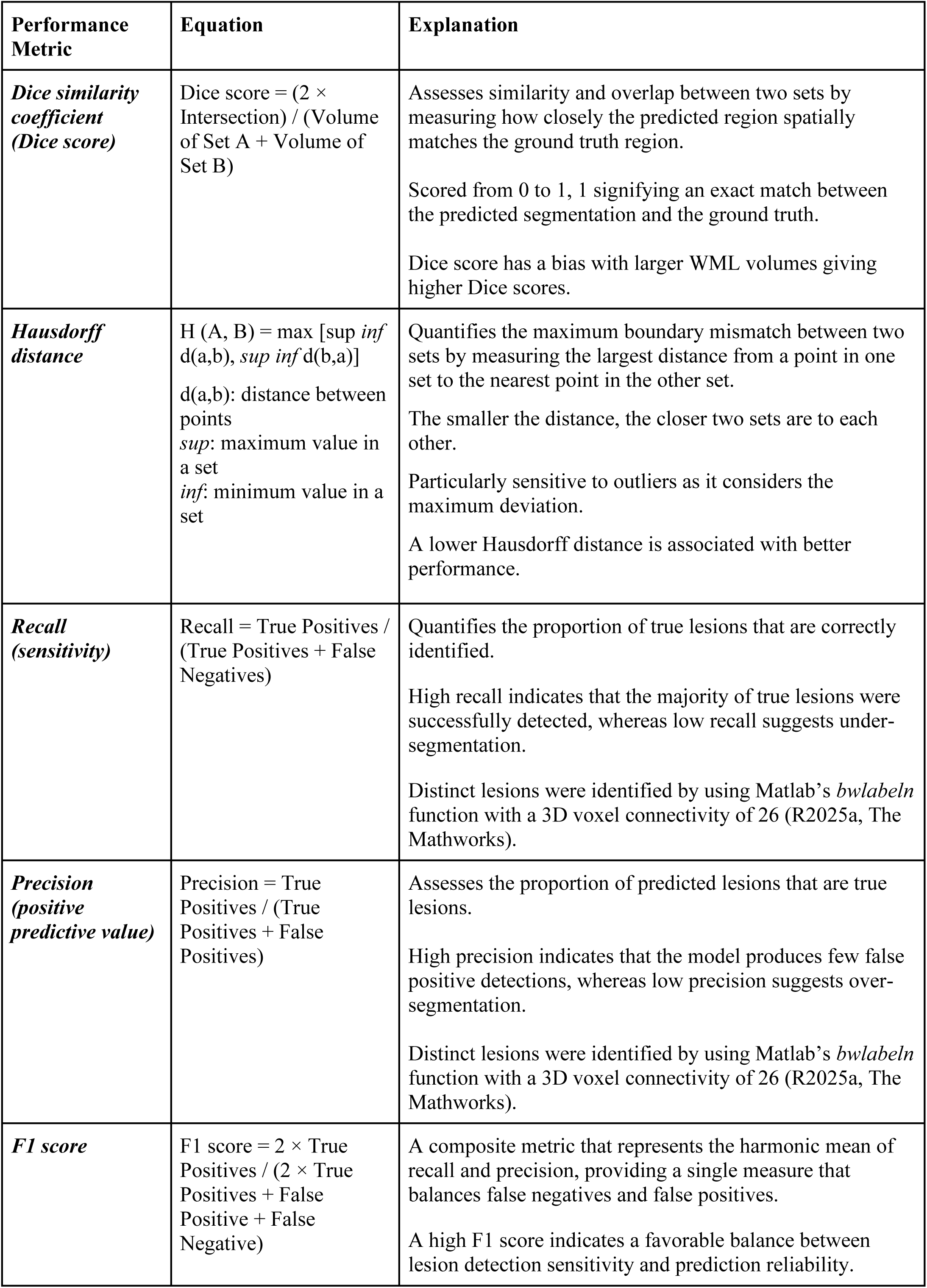
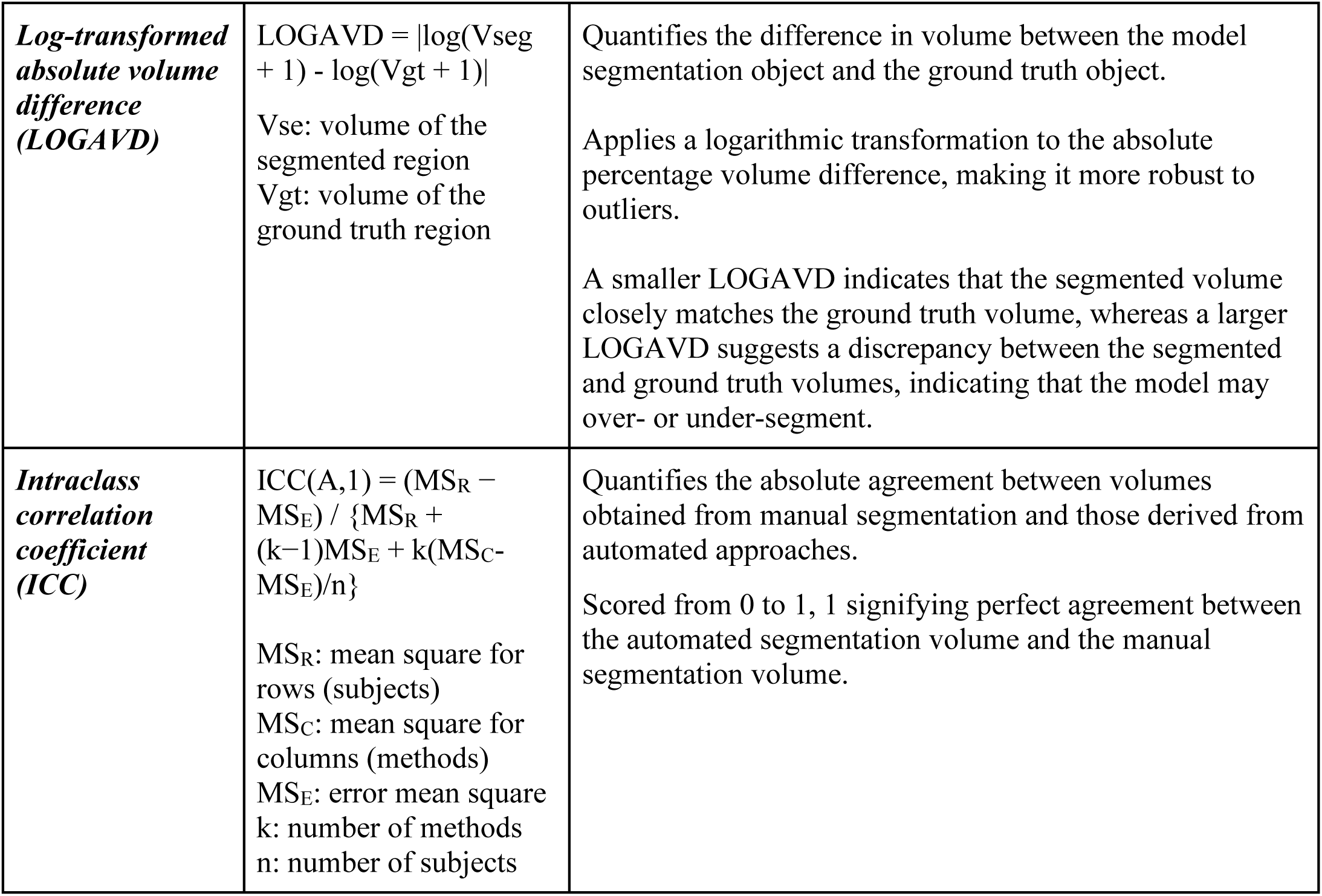
Summary of the metrics used to assess the performance of the automated segmentation approaches.

### Correlation with Age and Visual Rating Scales

We also investigated the correlation of WML measures with visual rating scales and age. Visual rating scales are used clinically to identify the location and severity of WML, albeit they do not quantify WML volume (Fazekas et al., 1987; Wahlund et al., 2001). The Fazekas and Wahlund scales are most commonly used for evaluating WML burden in clinical practice and trials and are therefore a clinically meaningful comparator for assessing the performance of automated approaches. In addition, age is one of the most important risk factors for WML, and it is well established that WML volume increases with age (regardless of other co-morbidities), so a positive correlation between age and WML burden would be expected (Griffanti et al., 2018; Hotz et al., 2022). This analysis was only performed within the PD group, as the majority of HC subjects were within the low lesion load group. PE conducted the visual ratings.

### Lobar Subgroup Analysis

We investigated whether the performance of each automated WML approach differed according to the WML location. For this, we measured WML distribution using predefined lobes: frontal, temporal, parietal, occipital, cingulate, insular lobe, and remaining unparcellated white matter corresponding with the centrum semiovale and corpus callosum categorized as deep white matter (DWM). Each lobe was parcellated using cortical-adjacent white matter parcellation by modifying the cortical parcellation provided by FreeSurfer (Desikan et al., 2006). Dice and ICC were calculated in each lobe.

### Visual Quality Control of WML Segmentations

In addition to quantitative metrics, we also ensured that all outputs from each of the automated approaches were visually inspected in ITK-SNAP (by SA). This was done to quality control the outputs, aiming to detect any systematic errors in the approaches we used and identify any patterns in under- or over-segmentation and missing lesions.

### Statistical Analysis

All statistical analysis was performed using IBM SPSS software version 25 (IBM Corporation, Armonk, NY, USA) and R software, v.4.0.3 (R Foundation for Statistical Computing, R-project.org). A p-value <0.05 was considered statistically significant.

#### Inter-rater Reliability of Manual Segmentations

Inter-rater reliability was assessed using Dice score and ICC on a subset of ten scans to ensure adequate agreement between the two raters.

#### Manual vs Automated Segmentations

Demographic features, clinical characteristics, and MRI scanner parameters were compared using independent t-tests and Pearson’s χ^2^-tests depending on the distribution of the data. We performed analysis of variance (ANOVA) tests for the evaluation metrics: Dice, Hausdorff, LOGAVD, recall, precision and F1 score compare the manual segmentation with the output from each automated approach. Variance was assessed using the Bartlett test (Bartlett, 1937); when variance was equal the Tukey’s HSD test was used (Tukey, 1949), when unequal the Games Howell test was used for post-hoc analyses to identify which pairs of means significantly differed (Games and Howell, 1976). ICC was calculated using the Reliability Analysis procedure in IBM SPSS (version 25; IBM Corporation, Armonk, NY, USA), specifying a two-way mixed-effects model, single measurement, and absolute agreement definition [ICC(A,1)], which is appropriate when the algorithms are treated as fixed effects and the objective is to evaluate their absolute agreement rather than consistency alone and account for systematic over- or under-estimation by an algorithm.

#### Correlation with Age and Visual Rating Scales

Bivariate correlation analysis was performed to assess correlation between age and WML volumes from the manual segmentations and automated approaches. Partial correlation analysis after adjusting for age was used to form correlations with visual rating scales (Fazekas and Wahlund). Family-wise error rate corrected p-value of <0.05 using Bonferroni correction was applied for multiple comparisons.

## RESULTS

### Demographic and Clinical Characteristics of Participants

The demographic and clinical features of the PD and HC groups are summarized in **Table 3**. Data were collected from 204 PD and 72 HC; after exclusion of 3 PD and 8 HC participants due to segmentation failures, 201 PD and 64 HC subjects were included in the analysis. In the PD group, the mean age was 66.57 ± 7.86 years with 34.8% female. In the HC group, the mean age was 66.26 ± 8.67 with 43.8% female. Both age and sex did not significantly differ between the two groups. In the PD group, the mean Unified Parkinson’s Disease Rating Scale (UPDRS) total score was 34.30 ± 14.74 and the Hoehn and Yahr scale was 1.92 ± 0.68. The mean Montreal Cognitive Assessment (MoCA) score significantly differed between the two groups: in the PD group, the average score was slightly lower, with 26.33 ± 3.3, compared to 28.07 ± 1.17 in the HC group. However, both means were within the normal cognitive range. The purpose of this study was not to study clinical correlations with WML, but to identify an optimal automated segmentation approach; however, detailed demographic features are provided in **Table 3** to highlight disease variability. All scanner parameters are outlined in **Table 4**.

**Table 3:** Demographic features, clinical characteristics, and scanner parameters for participants. Values are expressed as mean ± standard deviation or number (percentage). P-values are the results of independent t-tests and chi-squared tests, as appropriate. Abbreviations: MDS-UPDRS, The Movement Disorder Society-Unified Parkinson’s Disease Rating Scale; MoCA, Montreal Cognitive Assessment; PD, Parkinson’s disease; WML, white matter lesions.

|  | Parkinson's Disease | Control | p-value |
| --- | --- | --- | --- |
| Number | 201 | 64 | - |
| Lesion load |  |  | **0.005** |
| Low lesion load (%) | 145 (72.1) | 57 (89.1) |  |
| Medium lesion load (%) | 39 (19.4) | 3 (4.7) |  |
| High lesion load (%) | 17 (8.5) | 4 (6.2) |  |
| Age, yr | 66.57 ± 7.86 (n = 201) | 66.26 ± 8.67 (n = 64) | 0.802 |
| Low WML Load | 64.91 ± 7.56 (n = 145) | 66.23 ± 8.82 (n = 57) | 0.321 |
| Medium WML Load | 69.38 ± 6.71 (n = 39) | 65.50 ± 7.03 (n = 3) | 0.442 |
| High WML Load | 74.31 ± 6.48 (n = 17) | 67.30 ± 9.42 (n = 4) | 0.089 |
| Sex, female (%) | 70 (34.8) (n = 201) | 28 (43.8) (n = 64) | 0.255 |
| Low WML Load | 48 (33.1) (n = 145) | 24 (42.1) (n = 57) | 0.229 |
| Medium WML Load | 16 (41.0) (n = 39) | 1 (33.3) (n = 3) | 1.000 |
| High WML Load | 6 (35.3) (n = 17) | 3 (75) (n = 4) | 0.272 |
| MDS-UPDRS total score | 34.30 ± 14.74 (n = 135) | - | - |
| Low WML Load | 33.36 ± 14.32 (n = 104) | - | - |
| Medium WML Load | 36.68 ± 13.43 (n = 22) | - | - |
| High WML Load | 39.44 ± 21.65 (n = 9) | - | - |
| Hoehn and Yahr scale | 1.92 ± 0.68 (n = 135) | - | - |
| Low WML Load | 1.87 ± 0.68 (n = 104) | - | - |
| Medium WML Load | 1.95 ± 0.58 (n = 22) | - | - |
| High WML Load | 2.44 ± 0.73 (n = 9) | - | - |
| MoCA | 26.33 ± 3.3 (n = 135) | 28.07 ± 1.17 (n = 44) | **< 0.001** |
| Low WML Load | 26.48 ± 3.27 (n = 104) | 28.21 ± 1.17 (n = 39) | **< 0.001** |
| Medium WML Load | 26.27 ± 2.91 (n = 22) | 27.00 ± 0.00 (n = 3) | 0.255 |
| High WML Load | 24.67 ± 4.36 (n = 9) | 27.00 ± 0.00 (n = 2) | 0.147 |

**Table 4:** Summary of MRI scanner parameters. Data presented for the entire Parkinson’s disease and control groups and lesion load subgroups where applicable. TE = Echo Time; TR = Repetition Time; TI = Inversion Time; 2D = Two-dimensional; 3D = Three-dimensional

| Parameter | Parkinson's Disease | Control |
| --- | --- | --- |
| Sample Size | 201 | 64 |
| Data Source | C-BIG: 48 (23.9%);<br>PPMI: 96 (47.8%);<br>UPenn: 57 (28.4%) | C-BIG: 7 (10.9%);<br>PPMI: 44 (68.8%);<br>UPenn: 13 (20.3%) |
| *Low WML Load* | C-BIG: 29 (20.0%);<br>PPMI: 79 (54.5%);<br>UPenn: 37 (25.5%) | C-BIG: 6 (10.5%);<br>PPMI: 39 (68.4%);<br>UPenn: 12 (21.1%) |
| *Medium WML Load* | C-BIG: 11 (28.2%);<br>PPMI: 14 (35.9%);<br>UPenn: 14 (35.9%) | C-BIG: 0 (0.0%);<br>PPMI: 3 (100.0%);<br>UPenn: 0 (0.0%) |
| *High WML Load* | C-BIG: 8 (47.1%);<br>PPMI: 3 (17.6%);<br>UPenn: 6 (35.3%) | C-BIG: 1 (25.0%);<br>PPMI: 2 (50.0%);<br>UPenn: 1 (25.0%) |
| MRI Field Strength | 3T: 140 (69.7%); 1.5T: 61 (30.3%) | 3T: 37 (57.8%); 1.5T: 27 (42.2%) |
| *Low WML Load* | 3T: 95 (65.5%); 1.5T: 50 (34.5%) | 3T: 32 (56.1%); 1.5T: 25 (43.9%) |
| *Medium WML Load* | 3T: 30 (76.9%); 1.5T: 9 (23.1%) | 3T: 2 (66.7%); 1.5T: 1 (33.3%) |
| *High WML Load* | 3T: 15 (88.2%); 1.5T: 2 (11.8%) | 3T: 3 (75%); 1.5T: 1 (25%) |
| Scanner Manufacturer | GE: 55 (27.4%);<br>Philips: 18 (9%);<br>Siemens: 128 (63.7%) | GE: 25 (39.1%);<br>Philips: 11 (17.2%);<br>Siemens: 28 (43.8%) |
| *Low WML Load* | GE: 46 (31.7%);<br>Philips: 15 (10.3%);<br>Siemens: 84 (57.9%) | GE: 21 (36.8%);<br>Philips: 10 (17.5%);<br>Siemens: 26 (45.6%) |
| *Medium WML Load* | GE: 8 (20.5%);<br>Philips: 2 (5.1%);<br>Siemens: 29 (74.4%) | GE: 2 (66.7%);<br>Philips: 1 (33.3%);<br>Siemens: 0 (%) |
| *High WML Load* | GE: 1 (5.9%);<br>Phillips: 1 (5.9%);<br>Siemens: 15 (88.2%) | GE: 2 (50.0%);<br>Phillips: 0 (0.0%);<br>Siemens: 2 (50.0%) |
| **T1-weighted** |  |  |
| Acquisition Type | 3D: 201 (100%) | 3D: 64 (100%) |
| Slice Thickness (mm) | 1.0 - 2.0 | 1.0 - 2.0 |
| Voxel Size (mm) | [0.43-1.51] x [0.43-1.25] x [0.72-4.00] | [0.43-2.00] x [0.43-1.25] x [0.49-2.00] |
| TE (s) | 0.002 - 0.009 | 0.003 - 0.005 |
| TR (s) | 0.007 - 2.400 | 0.007 - 2.400 |
| TI (s) (where applicable) | 0.04 - 1.1 | 0.40 - 1.1 |
| Flip Angle (°) | 8 - 90 | 8 - 20 |
| Matrix Size | [84-512] × [192-512] × [34-320] | [80-512] × [192-512] × [72-512] |
| **FLAIR** |  |  |
| Acquisition Type | 2D: 144 (71.6%) / 3D: 57 (28.4%) | 2D: 51 (79.7%) / 3D: 13 (20.3%) |
| *Low WML Load* | 2D: 108 (74.5%) / 3D: 37 (25.5%) | 2D: 45 (78.9%) / 3D: 12 (21.1%) |
| *Medium WML Load* | 2D: 25 (64.1%) / 3D: 14 (35.9%) | 2D: 3 (100.0%) / 3D: 0 (0.0%) |
| *High WML Load* | 2D: 11 (64.7%) / 3D: 6 (35.3%) | 2D: 3 (75.0%) / 3D: 1 (25.0%) |
| Slice Thickness (mm) | 1.0 - 6.0 | 1.0 - 5.0 |
| Voxel Size (mm) | [0.45-1.05] x [0.43-1.05] x [1.00-7.47] | [0.43-1.00] x [0.43-1.05] x [1.00-8.47] |
| TE (s) | 0.074 - 0.292 | 0.100 - 0.292 |
| TR (s) | 6.00 - 11.00 | 6.00 - 12.65 |
| TI (s) | 1.65 - 2.80 | 2.0 - 2.8 |
| Flip Angle (°) | 90 - 180 | 90 - 180 |
| Matrix Size | [160-576] × [256-576] × [20-256] | [160-560] × [256-560] × [18-256] |

In the PD group, 72.1% had low lesion load, 19.4% had medium lesion load, and 8.5% had high lesion load. The proportion of low, medium, and high lesion load in the HC group was 89.1%, 4.7%, and 6.2% respectively. As most automated approaches perform better in higher lesion loads (Gaubert et al., 2023), a predominance of low lesion load was selected based on visual inspection of the FLAIR scans.

### Inter-rater Reliability of Manual Segmentation

Of the ten random samples, the average WML volume was 1.694 (SD 1.602) cm^3^, with nine subjects in the low WML load group and one in the medium WML load group. Adequate agreement between the two raters was confirmed by an ICC of 0.974 (95% CI 0.900 – 0.994) with p-value < 0.001 and a good mean Dice score for a low lesion load of 0.424 (SD 0.170).

### Comparison of Performance Metrics Among Automated Approaches

The results for the performance metrics in the PD and HC groups are summarized in **Table 5** and **Figures 1 and 2**. In PD, U-Net achieved the highest Dice score (0.46 ± 0.21), recall (0.76 ± 0.25), precision (0.49 ± 0.25), F1 score (0.54 ± 0.22) and ICC (0.965). Post-hoc analyses showed all metrics were statistically significantly higher than all other automated approaches, except for the Dice score with BIANCA-LOO (0.9). In HC, U-Net also achieved the highest Dice score (0.39 ± 0.21), recall (0.62 ± 0.31), precision (0.63 ± 0.27), F1 score (0.56 ± 0.22) and ICC (0.967). Post-hoc analyses showed all metrics were statistically significantly higher than all other automated approaches, except for the Dice score with BIANCA-LOO (0.9) and LPA-FAST (0.4).

**Table 5:**
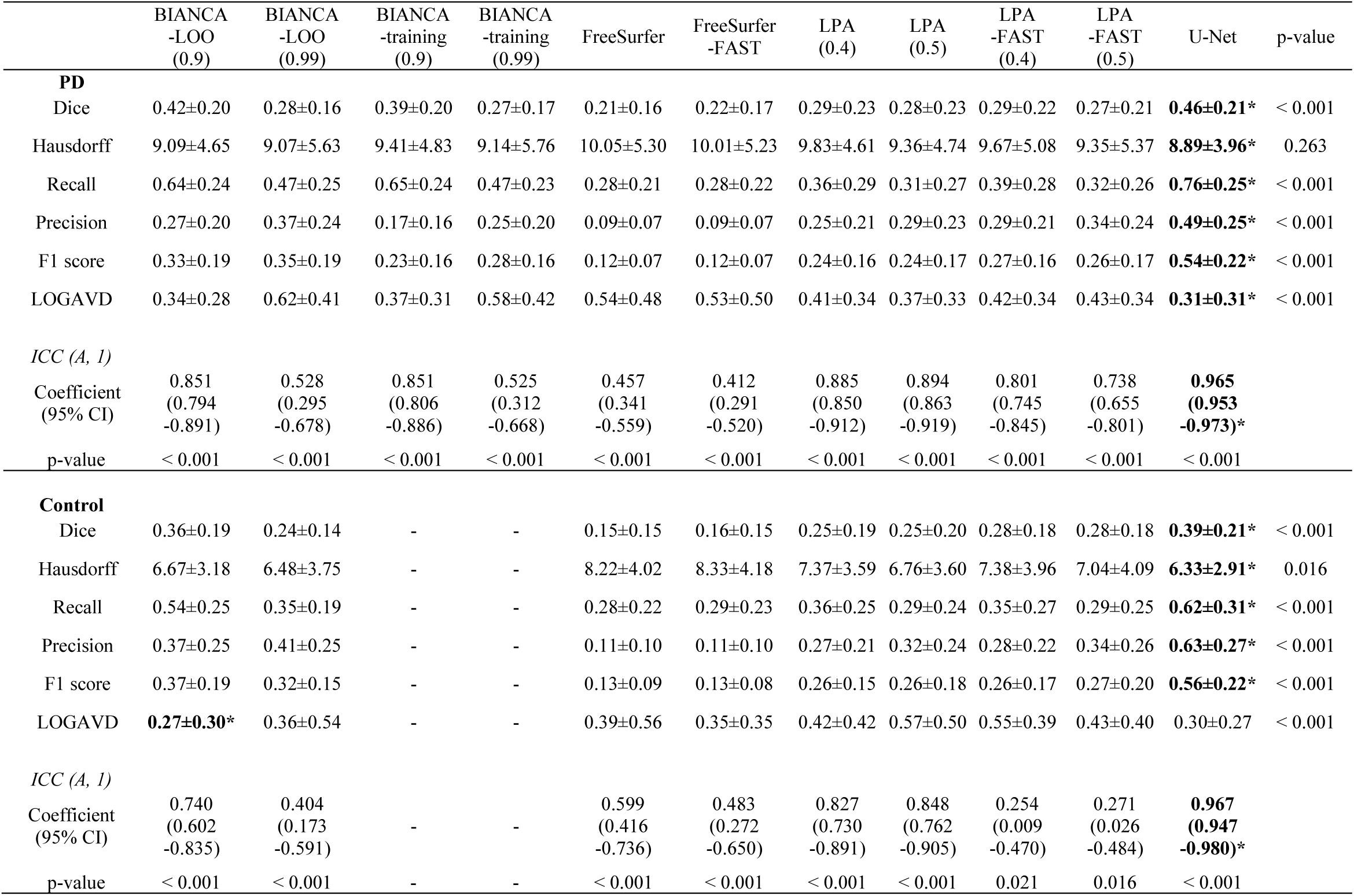
Performance metrics across the automated approaches in the PD and control group. Values in the upper part of the table are p-values of analysis of variance. The lower part of the table indicates intraclass correlation coefficients and their p-values. Numbers in parentheses indicate the thresholds used in the automated algorithms. The asterisk indicates the best value.

**Figure 1:**
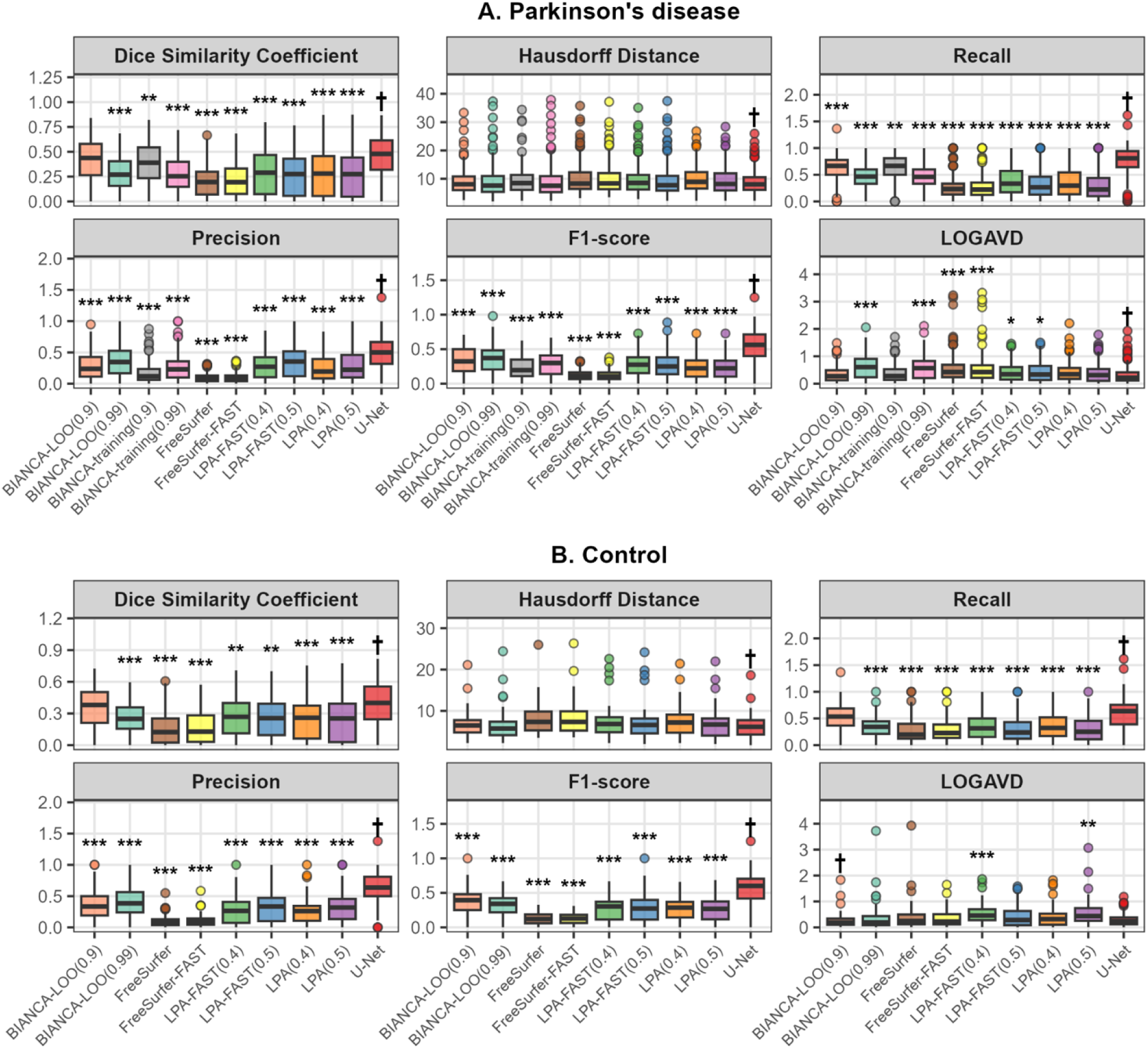
Boxplots of performance metrics across the automated approaches in the Parkinson’s disease group and control group. Comparison of white matter lesion segmentation performance across algorithms in Parkinson’s disease and control groups. Box plots show the distribution of performance metrics between automated segmentation algorithms and manual segmentation: (A) Parkinson’s disease group and (B) healthy controls. Each panel displays Dice Similarity Coefficient (higher values indicate better agreement), Hausdorff Distance (lower values indicate better agreement), recall (higher values indicate better agreement), precision (higher values indicate better agreement), F1 score (higher values indicate better agreement), and Log-transformed Absolute Volume Difference (LOGAVD; lower values indicate better agreement). The daggers indicate the best-performing algorithms within each metric and group. The asterisks indicate statistically significant differences compared to the best-performer (*p < 0.05, **p < 0.01, ***p < 0.001; Tukey HSD test for homogeneous variances or Games-Howell test for heterogeneous variances following Bartlett’s test).

**Figure 2:**
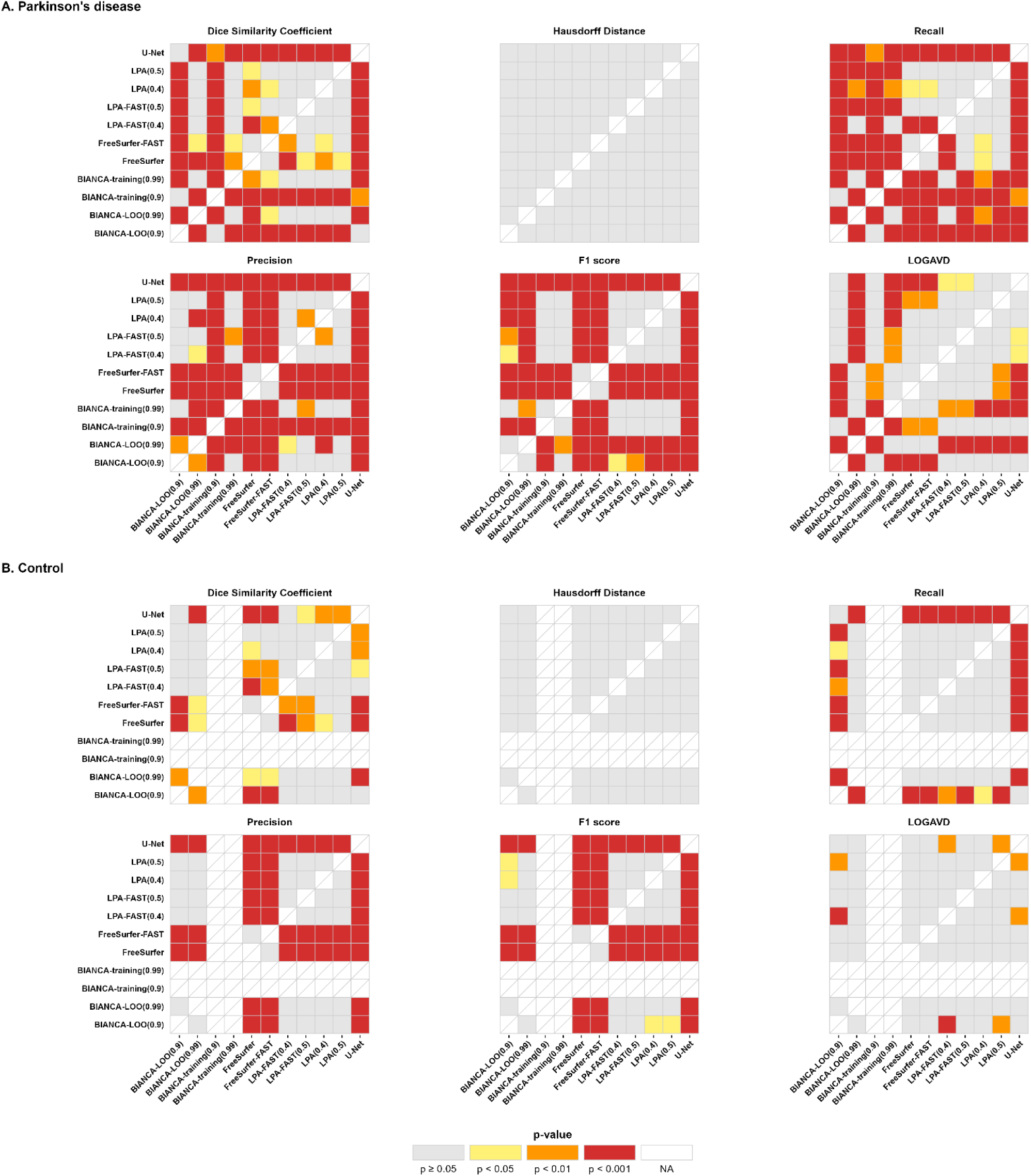
Post-hoc analysis of performance metrics across the automated approaches in the PD and control group. This heatmap illustrates the pairwise post-hoc comparisons of automated white matter lesion segmentation algorithms in Parkinson’s disease and control groups. Each cell displays the statistical significance (p-value) of differences in segmentation performance between two algorithms based on six evaluation metrics: Dice similarity coefficient, Hausdorff distance, recall, precision, F1 score, and log-transformed absolute volume difference (LOGAVD). Color intensity encodes the level of significance, with red indicating greater statistical difference and white indicating non-significant difference. Pairwise comparisons were conducted using either Games–Howell or Tukey’s HSD test, depending on the equality of variance across groups.

LOGAVD was lowest (i.e., best) for U-Net in PD (0.31 ± 0.31) and BIANCA-LOO (0.9) in HC (0.27 ± 0.30); however, the differences between approaches were not statistically significant in post-hoc analyses. The Hausdorff distance was lowest (i.e., better) with U-Net in both the PD and the control groups, with mean values of 8.89 ± 3.96 and 6.33 ± 2.91, respectively. However, the differences were not statistically significant in the post-hoc analysis.

Bland-Altman plots for comparison of the automated algorithms with manual segmentation are shown in **Figure 3**, visualizing the agreement between the two methods by showing how the differences vary with increasing WML volume. The WML volumes were not normally distributed, so a non-parametric approach was taken using the median, 2.5th and 97.5th percentiles.

**Figure 3:**
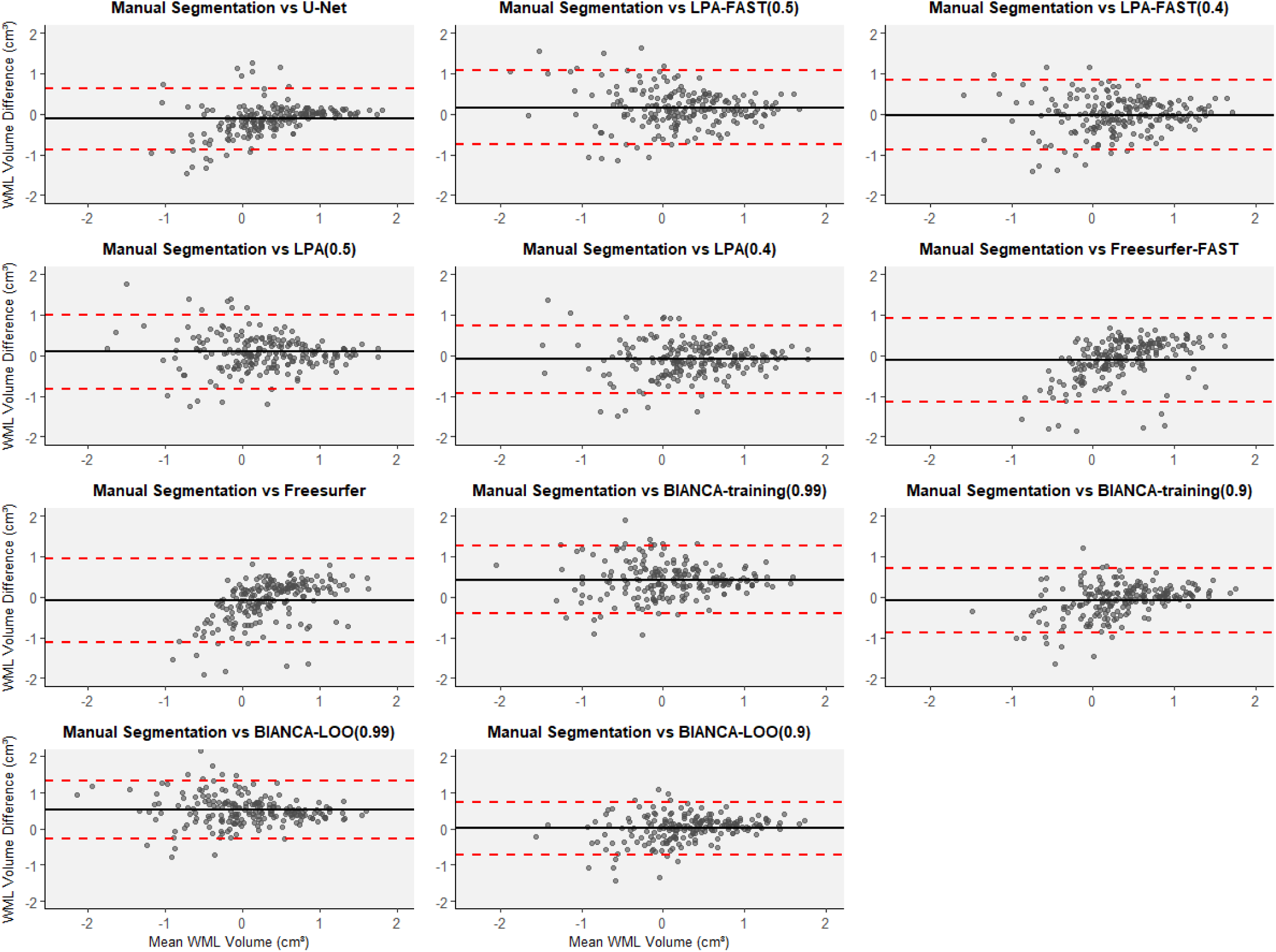
Bland-Altman plots for WML volume for the different algorithms in the PD group. Each plot illustrates the agreement between an automated algorithm and the manual segmentation by showing the difference in white matter lesion (WML) volume, calculated by subtracting the automated measurement from the manually segmented volume, plotted against the mean volume of the two methods across subjects. In this non-parametric Bland–Altman analysis, the central solid line represents the median of the differences, while the dashed lines indicate the 2.5th and 97.5th percentiles.

### Lesion Load Subgroup Analysis

#### Low WML Load Group

Results from the low WML load group are shown in **Table 6** and **Figure 4**. In PD, U-Net achieved the highest Dice score (0.39 ± 0.17), recall (0.74 ± 0.27), precision (0.44 ± 0.26), F1 score (0.49 ± 0.22) and ICC (0.569). Post-hoc analyses showed all metrics were statistically significantly higher for U-Net than for all other automated approaches, bar Dice with BIANCA-LOO (0.99) and recall with BIANCA-training (0.9) [**Figure 5**].

**Figure 4:**
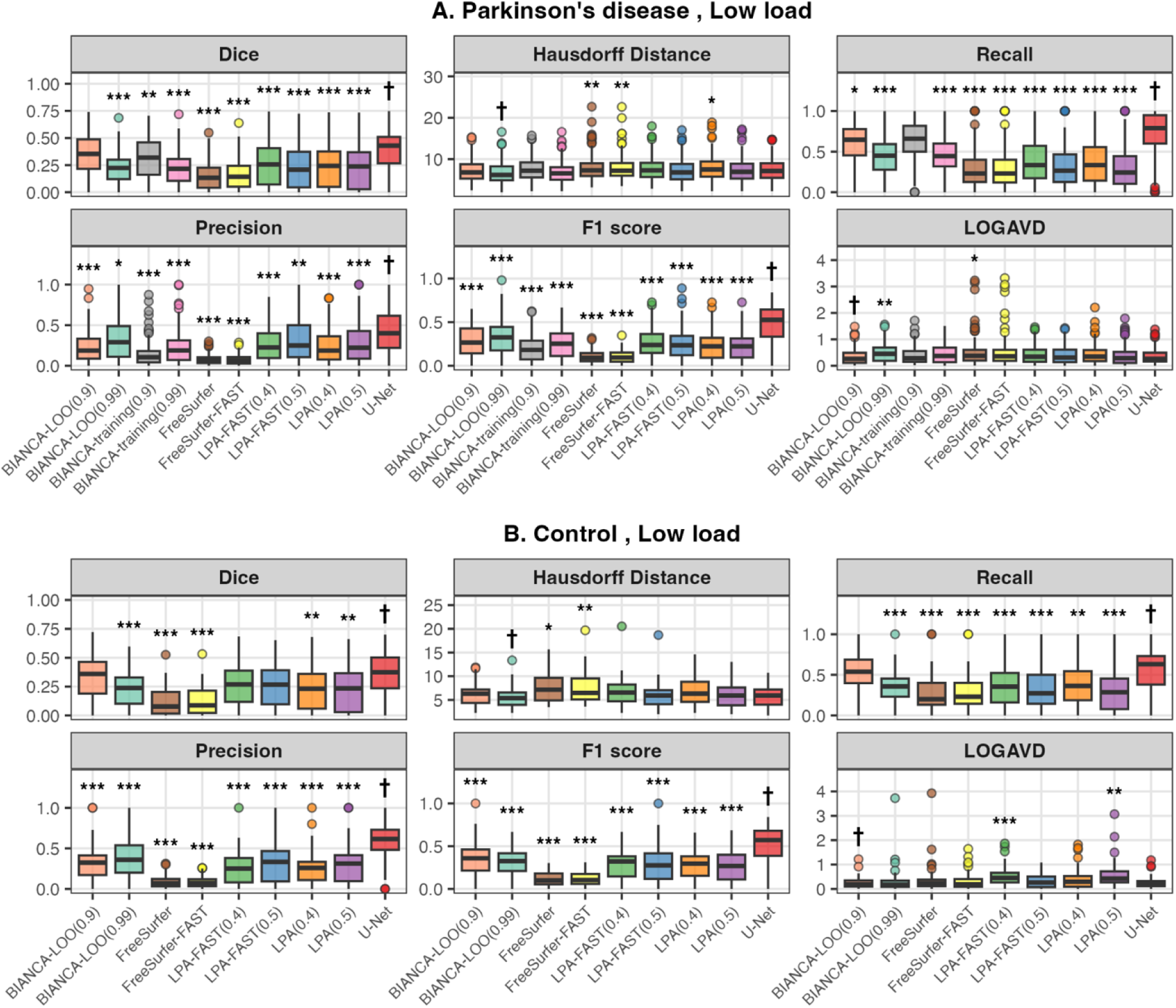
Boxplots of evaluation metrics across the automated approaches in the Parkinson’s disease group and control group for low white matter lesion load. Comparison of white matter lesion segmentation algorithm performance in subjects with low lesion load (volume <5 cm³). Performance of 11 automated segmentation algorithms was evaluated against manual segmentation using six agreement metrics: Dice Similarity Coefficient (higher values indicate better agreement), Hausdorff Distance (lower values indicate better agreement), recall (higher values indicate better agreement), precision (higher values indicate better agreement), F1 score (higher values indicate better agreement), and Log-transformed Absolute Volume Difference (LOGAVD; lower values indicate better agreement). (A) Parkinson’s disease patients with low lesion load, (B) healthy controls with low lesion load. Note that BIANCA-training algorithms were not applied to the control group. The daggers indicate the best-performing algorithms within each metric and group. The asterisks indicate statistically significant differences compared to the best-performer (*p < 0.05, **p < 0.01, ***p < 0.001; Tukey HSD test for homogeneous variances or Games-Howell test for heterogeneous variances following Bartlett’s test).

**Figure 5:**
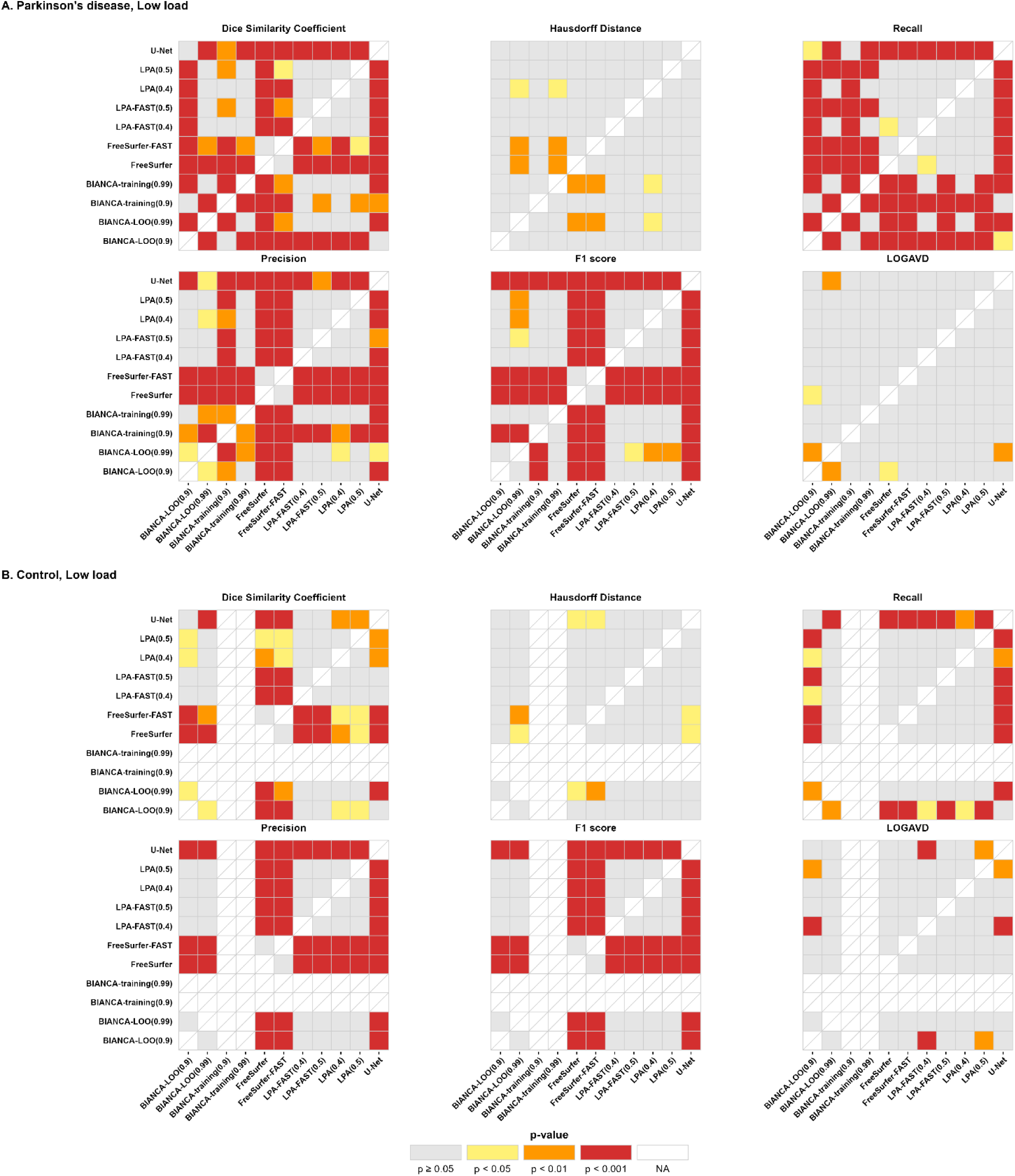
Post-hoc analysis of performance metrics across the automated approaches in the Parkinson’s disease group and control group for low white matter lesion load. This figure presents heatmaps of pairwise post-hoc comparisons of automated white matter lesion segmentation algorithms in subjects with low lesion load (volume <5 cm³). Panel A corresponds to Parkinson’s disease patients with low lesion load, while panel B displays results from healthy controls with low lesion load. Note that BIANCA-training algorithms were not applied to the control group. Each cell displays the statistical significance (p-value) of differences in segmentation performance between two algorithms based on six evaluation metrics: Dice similarity coefficient, Hausdorff distance, recall, precision, F1 score, and log-transformed absolute volume difference (LOGAVD). Red indicates greater statistical difference, and white indicates non-significant difference. Post-hoc comparisons were conducted using either Games–Howell or Tukey’s HSD test, depending on the equality of variance.

**Table 6:**
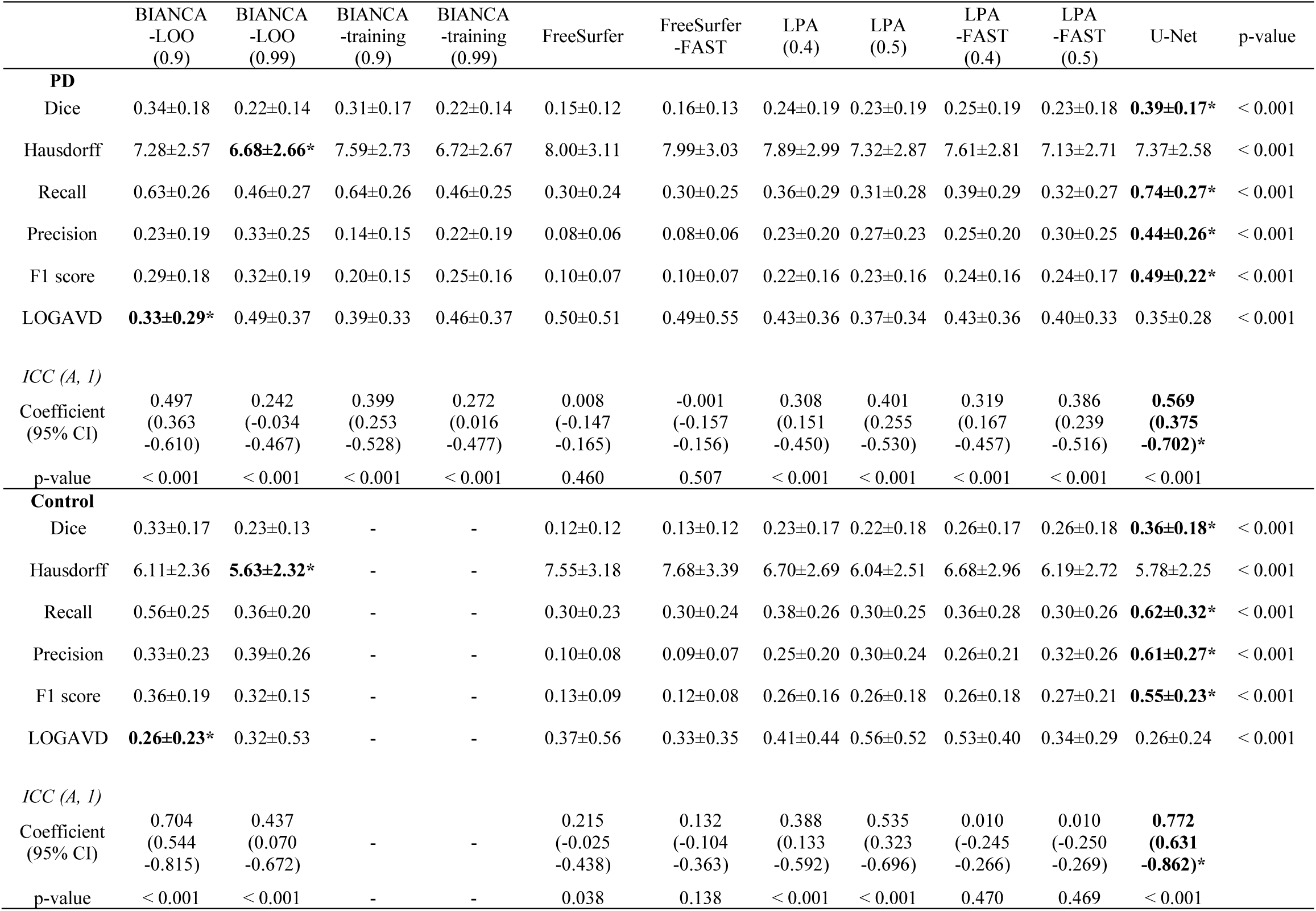
Performance metrics across the automated approaches in the PD and control group with low WML loads. Values in the upper part of the table are p-values of analysis of variance. The lower part of the table indicates intraclass correlation coefficients and their p-values. Numbers in parentheses indicate the threshold used in the automated algorithms. The asterisk indicates the best value.

In HC, U-Net also had the highest Dice score (0.36 ± 0.18), but the difference was not significant with BIANCA-LOO (0.9) and LPA-FAST (0.4) and (0.5). U-Net also showed the highest recall (0.62 ± 0.32), precision (0.61 ± 0.27), F1 score (0.55 ± 0.23) and ICC (0.772). Post-hoc showed all metrics were significantly higher for U-Net than for all other automated approaches except recall in BIANCA-LOO (0.9) [**Figure 5**].

BIANCA-LOO (0.9) achieved the lowest LOGAVD scores in both PD (0.33 ± 0.29) and HC (0.26 ± 0.23). The LOGAVD values only reached statistical significance in the PD group with BIANCA-LOO (0.99) and FreeSurfer and in the HC group with LPA-FAST (0.4) and LPA (0.5) [**Figure 5**]. In both PD and HC, BIANCA-LOO (0.99) achieved the lowest Hausdorff distance (PD: 6.68 ± 2.66, HC: 5.63 ± 2.32). However, this was not statistically significant in the PD group but was in HC when compared with LPA (0.4), FreeSurfer and FreeSurfer-FAST [**Figure 5**].

#### Medium WML Load Group

Results from the medium WML load group are shown in **Supplementary Table 1** and **Supplementary Figure 1.** In PD, U-Net achieved the highest Dice score (0.61 ± 0.15), recall (0.76 ± 0.21), precision (0.61 ± 0.19), F1 score (0.64 ± 0.16) and ICC (0.583). Post-hoc analyses showed all metrics for U-Net were statistically significantly higher than for all other automated approaches, bar Dice with BIANCA-LOO (0.9) and recall with BIANCA-LOO (0.9) and BIANCA-training (0.9) [**Supplementary Figure 2**]. U-Net also achieved the lowest Hausdorff distance (12.31 ± 3.97); however, this did not significantly differ from any of the other automated approaches. LOGAVD was also lowest with U-Net (0.25 ± 0.42); this value was significantly different to BIANCA-LOO (0.99), BIANCA-training (0.99), FreeSurfer, and FreeSurfer-FAST [**Supplementary Figure 2**].

#### High WML Load Group

Results from the high WML load group are shown in **Supplementary Table 2** and **Supplementary Figure 3.** In PD, U-Net achieved the highest Dice score (0.77 ± 0.07), recall (0.85 ± 0.10), precision (0.64 ± 0.13), F1 score (0.72 ± 0.10) and ICC (0.941). Post-hoc analyses showed all metrics for U-Net were significantly higher than for all other automated approaches except for Dice with BIANCA-LOO (0.9) and BIANCA-training (0.9) and precision with BIANCA-LOO (0.99) [**Supplementary Figure 4**]. U-Net achieved the lowest Hausdorff distance (14.00 ± 4.58), but the difference between other approaches was not significant in post-hoc analysis. LOGAVD score was lowest with U-Net (0.14 ± 0.09), but the difference was not significant with BIANCA-training (0.9), LPA-FAST (0.4), LPA (0.4) and LPA (0.5) [**Supplementary Figure 4].**

### Lobar Subgroup Analysis

We performed subgroup analysis according to frontal, temporal, parietal, occipital, cingulate, insular lobes, and deep white matter. The results are presented in **Table 7** and **Supplementary Table 3**. In both groups, the Dice score in U-Net was highest in all lobes and deep white matter. U-Net had the highest ICC in all regions in PD and all except the occipital region in controls, where LPA-FAST (0.5) was highest.

**Table 7:**
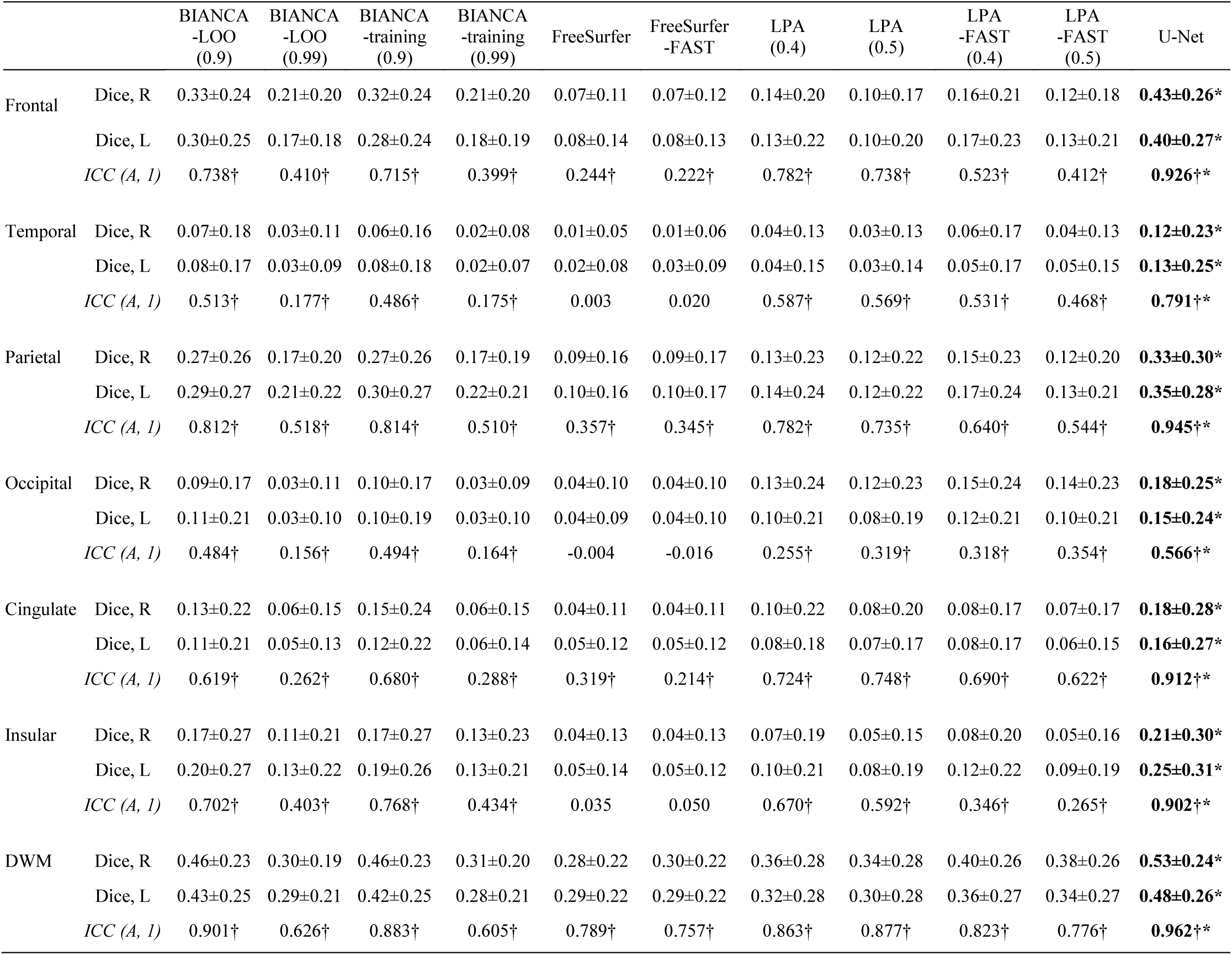
Dice score and ICC of automated WML measuring approaches according to lobes in the PD group. Values are expressed as mean ± standard deviation or number (percentage). The dagger indicates statistical significance. The asterisk indicates the best value.

### Correlation with Age and Visual Rating Scales

#### Correlation of Outputs Using Age

In the PD group, WML volumes from all the automated segmentation approaches were correlated with age [**Supplementary Table 4**]. LPA (0.5) had the highest Pearson correlation coefficient (r = 0.489, p < 0.001), which was higher than that of manual segmentation (r = 0.434, p < 0.001).

#### Correlation of Outputs Using Visual Rating Scales

In the PD group, the WML volumes from the manual segmentation and from all the automated segmentation approaches were well correlated to both Fazekas and Wahlund score, after adjusting for age [**Supplementary Table 5**]. U-Net had the highest partial correlation coefficient among the automated approaches (Fazekas r = 0.805, p < 0.001, Wahlund r = 0.811, p < 0.001) with similar values to that of the manual segmentation (Fazekas r = 0.863, p < 0.001, Wahlund r = 0.874, p < 0.001).

### Visual Quality Control of WML Segmentations

All outputs from the automated approaches were visually compared to the manual segmentations. U-Net and BIANCA tended to produce the most accurate maps. However, small deep white matter lesions were often missed with the BIANCA approach after thresholding. U-Net was noted to detect both periventricular and deep white matter lesions well but had a tendency to identify the optic radiation as WML [**Figure 6**]. All approaches produced errors in a proportion of the segmentation outputs, but we failed to identify any particular pattern in the imaging parameters which led to these errors, emphasizing the need for quality checking of all outputs, as no automated approach is foolproof.

**Figure 6:**
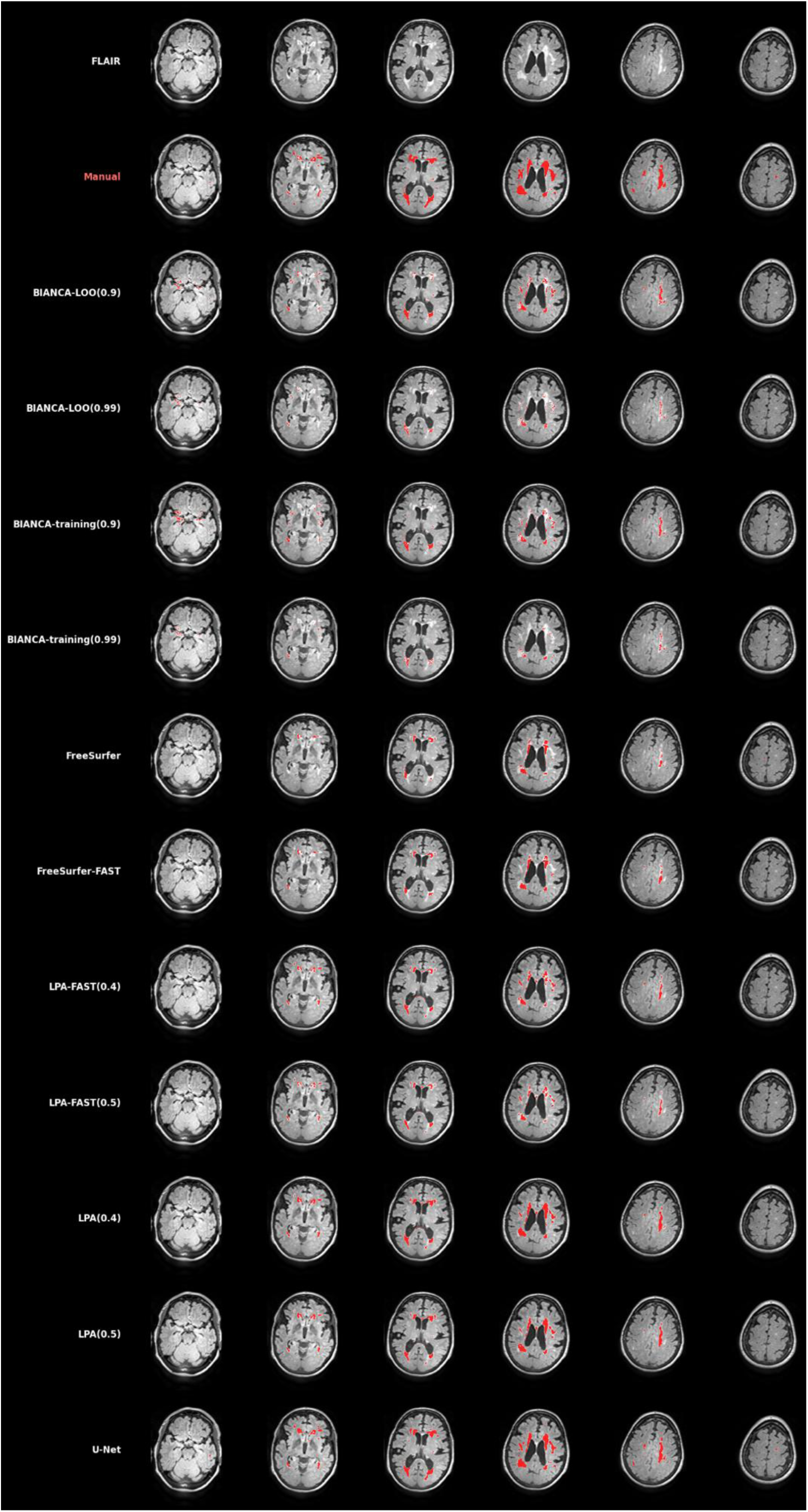
An example segmentation from each of the automated approaches. Top row shows FLAIR MRI slices from a single example subject. The segmentation provided by each automated approach is shown in subsequent rows.

## DISCUSSION

The purpose of this study was to compare and identify an optimal WML automated segmentation approach to use on multi-site data in PD, which includes data from various scanners and acquisition parameters. Of those assessed, U-Net was the best-performing automated approach overall relative to the gold standard of manual segmentation, reflected by the overwhelming majority of comparisons showing most favorable Dice and ICC values for spatial and volumetric agreement and consistently more favorable F1, recall and precision scores, reflecting a good balance of true positives and false negatives. This pattern was seen consistently across various scanner parameters in both PD and HC.

U-Net consistently outperformed all other approaches in the full PD and HC analysis; however, when sub-grouped into lesion loads, there were some occasions where other approaches performed best. In the low WML load group, Hausdorff distance was lowest in BIANCA-LOO (0.99) and LOGAVD in BIANCA-LOO (0.9) for both PD and HC. As Hausdorff distance is highly sensitive to outliers (Taha and Hanbury, 2015), even a single outlier can substantially impact the calculated distance. As many of our scans had dot-like small lesions, this might have led to a misleading comparison between approaches in the post-hoc analysis. Additionally, overestimation of the optic radiation with U-Net, which was identified in visual quality control, may account for the poorer performance on LOGAVD. Even a small absolute difference in volume can substantially impact LOGAVD score, as this logarithmic metric compresses large differences and expands small ones, potentially exaggerating the perceived error. Hence it is important to combine various performance metrics to not only understand volumetric and spatial agreement, but what drives them.

Although BIANCA-LOO performed best on certain metrics discussed above, the BIANCA-LOO approach is not scalable in multi-site studies as it requires manual labelling of all subjects to create subject-specific training sets. Instead, the BIANCA-LOO analysis can be interpreted as an approximate upper bound on the performance achievable by BIANCA when all available manually labelled data are used for training. In practice, BIANCA’s performance depends strongly on the quality and representativeness of the training set. Future work should investigate the impact of larger and more heterogeneous training datasets.

In the lobar subgroup analysis, U-Net also had the highest Dice and ICC across all lobes in the PD group and controls, except the occipital lobe in controls for which the ICC was highest for LPA-FAST (0.5). The tendency of U-Net to misclassify the optic radiation as WML may explain its relatively weaker performance in the occipital lobe. Studies assessing WML in PD have produced conflicting results which may be attributable to differences in segmentation approach, for example under-identifying lesions in clinically eloquent brain regions (Gattellaro et al., 2009). Therefore, an approach that accurately detects the location of WML, regardless of which lobe lesions are located in, may reduce the risk of missing clinically relevant lesions in PD.

Our findings in PD are largely consistent with current literature on the topic in other populations. The MICCAI WMH Segmentation Challenge aimed to compare the segmentation of WML of presumed vascular origin on MR images using automated approaches in patients from five memory and aging cohorts. Results revealed no single method excelled in all metrics, yet U-Net generally performed better at various lesion loads (Kuijf et al., 2019). Results also showed that ensemble methods (such as U-Net, which we implemented) outperformed other convolutional neural networks (CNN), highlighting that smaller WML were often missed by the other CNN methods and methods that were less ‘location sensitive’ were better in this regard (Ghafoorian et al., 2016; Kuijf et al., 2019). As both the size and shape of WML are clinically impactful, it is essential to select an approach that does not miss these smaller lesions (De Bresser et al., 2018). Torres-Simon et al. demonstrated in healthy aging participants that although LPA and BIANCA outperformed other approaches tested (they did not include U-Net in their comparison), they failed to consistently segment lesions smaller than 8 mm^3^. They argued deep learning approaches show promise for small lesion segmentation due to the utility of multiscale deep features (Park et al., 2021; Torres-Simon et al., 2024). Studies comparing BIANCA and other k-nearest neighbor techniques in healthy aging and cognitively impaired populations concluded BIANCA/k-NN-TTP followed by LST-LPA outperformed other methods tested, including in longitudinal data, while FreeSurfer consistently underestimated WML volumes (Heinen et al., 2019; Ribaldi et al., 2021; Hotz et al., 2022). Interestingly, a recent review of WML automated approaches found no evidence that deep learning was superior to other supervised or unsupervised techniques; however, this discrepancy likely arises as the inclusion criteria encompassed both WML of vascular origin and inflammatory WML such as seen in multiple sclerosis. Most of the studies included in the review did not use multi-site data (Balakrishnan et al., 2021).

In this study, we included data from various scanners and acquisition parameters to reflect consortia data which is often variable. U-Net performed well despite these differences. In addition, for an automated approach to be amenable to consortium data there are several logistical considerations. Various sites will have differing computational facilities and skillsets. Ideally, BIANCA requires a training set that is representative of the population, hence a training set may require combining labelled samples from different sites, increasing the level of manual input required compared to U-Net. Nonetheless, U-Net is computationally demanding and has a longer running time than BIANCA. Additionally, all automated approaches require the installation of software packages, file preparation and pre-processing steps. Future work should consider a containerized approach to large multi-site analysis i.e., a pipeline that includes all the code, tools and dependencies required to output the WML segmentations.

The extent to which WML segmentations reflect meaningful pathological or clinical correlates remains uncertain and was beyond the scope of this study. The assumption made by the authors is that a technique that produces accurate WML segmentations in heterogeneous data would allow for clinical correlations to be elucidated in further large-scale analyses. Work is in progress on implementing this pipeline to other disease states with vascular pathology such as stroke with initial promising results.

There are also other limitations to our study. Its cross-sectional design precluded assessment of longitudinal consistency or progression sensitivity, which are important for evaluating the practical applicability of these methods in clinical and research settings. Some longitudinal studies have shown ‘outlier intervals’, where factors such as atrophy and co-morbidities affected outcomes of approaches such as BIANCA (Hotz et al., 2022). However, recent studies suggest deep learning techniques such as U-Net may actually help in modelling longitudinal multi-modal MRI changes, including WML (Rachmadi et al., 2020; Gong et al., 2024). We also acknowledge that other automated segmentation approaches have been developed since this study was conducted. Tsuchida et al. (2023) found that their SHIVA-WMH tool produced more accurate segmentations than UNet-pgs; however, this comparison examined only 21 healthy subjects in two cohorts, with all scans collected on a 3T scanner with 1mm isotropic voxels (for T1w images) and 1.00 × 1.00 × 1.00 mm^3^ to 1.05 × 1.00 × 1.00 mm^3^ voxels (for FLAIR images). Additionally, UNet-pgs was used without retraining the model on the same training data used by SHIVA-WMH (Tsuchida et al., 2023). Finally, our study did not examine how individual scanner parameters affect accuracy of WML segmentations. Instead, we selected a heterogeneous sample based on a range of scanner and image acquisition parameters.

Our study also has several strengths. To our knowledge, it is the first attempt to evaluate automated WML algorithms specifically in PD, comparing their outputs to manual segmentations with whole brain and lobar assessments. Associations with age and visual rating scales were also made, allowing for comparison to alternative approaches. We used heterogeneous data with varying acquisition parameters and WML load and location, reflecting the real-world clinical environment and consortium data. We demonstrated promise for an approach that produces accurate WML segmentations despite these differing parameters across all brain regions, indicating U-Net’s potential utility for large, multi-site data analyses in PD and controls, with potential for detecting WML of vascular origin in other conditions.

## CONCLUSION

In summary, accurate and representative WML segmentations are key to analysis of the clinical impact of WML of presumed vascular origin. Optimization of an analysis approach on multi-site WML data specifically in a PD population was lacking. This paper outlines our attempts to select an optimal WML segmentation approach, in multi-site data, by comparing WML automated segmentation methods against the gold standard of manual segmentations. We found that U-Net has emerged as the best performing automated method for WML segmentation overall in PD and controls, consistently outperforming other algorithms across various lesion loads and in most lobar regions. Although BIANCA-LOO also performed to a high standard, this requires manual segmentation of all subjects in the dataset, which is not feasible for large-scale analyses. No automated approach was without limitations, therefore manual quality control of the outputs from all the automated approaches is essential. To expand the utility of U-Net, future work should focus on incorporating U-Net into a containerized imaging pipeline to streamline large-data analyses of WML patterns and validation of U-Net in other disease states.

## AUTHORSHIP

Conceptualization: SA, HH; Data curation: SA, SHY, PE, SMA, AA, AC, ES, EB, MA, SK, RPT, JB, COW, CTM, SIT, AD, HH; Formal analysis: SA, SHY, PE, SMA, JC, AA, ES, EB, MA, SK, RPT, LG, CV, HH; Funding acquisition: SA, HE, AS; Methodology: SA, SHY, SMA, JC, AA, LG, NJ, PMT, YVDW, CV, HE, LP, HH; Project administration: SA, SIT, AS, HH; Software: SHY, SMA, AA, JCK, LG, CV, HH; Supervision: SA, AS, HH; Visualization: SHY; Writing – original draft: SA, SHY, SMA, AS, HH; Writing – review & editing: all authors.

## DATA AVAILABILITY STATEMENT

All analysis code and derived data are available on request.

PPMI data is available at www.ppmi-info.org/data.

C-BIG data can be requested through the Quebec Parkinson Network Neuroimaging Cohort (https://zenodo.org/records/17450887) and approved upon reasonable request.

Restricted access data from UPenn can be requested through the Penn Neurodegeneration Data Sharing Committee (https://www.pennbindlab.com/data-sharing) and approved upon reasonable request.

## IRB STATEMENT

All participants gave informed consent for their participation and for sharing of their anonymized data from all three sources of data. Specifically for UPenn, all participants participated in an informed consent procedure approved by an Institutional Review Board (IRB) convened at the University of Pennsylvania and consented to participation and for sharing of their anonymized data. Regarding C-BIG data, the study was approved by the Research Ethics Board of the McGill University Health Centre.

Data used in the preparation of this article was obtained on [2021-03-01] from the Parkinson’s Progression Markers Initiative (PPMI) database (www.ppmi-info.org/access-data specimens/download-data), RRID:SCR_006431. For up-to-date information on the study, visit www.ppmi-info.org.

## Acknowledgements

We thank the ENIGMA-Parkinson’s Disease Working Group core team for guidance on study design, Ms Rida Waseem for statistical support, and Dr Nicola Rennie and Professor Joanne Knight for support based at Lancaster University.

We acknowledge the use of the UCL Myriad High Performance Computing Facility (Myriad@UCL), and associated support services, in the completion of this work. With special thanks to the Centre for Advanced Research Computing at UCL, the assistance given by Research IT and the use of the Computational Shared Facility at the University of Manchester, and the assistance and support of the IT team at the University of Lancaster.

Some data used in this article were obtained from the Parkinson’s Progression Markers Initiative (PPMI) database (www.ppmi-info.org/data). For up-to-date information on the study, visit www.ppmi-info.org. PPMI – a public-private partnership – is funded by the Michael J. Fox Foundation for Parkinson’s Research and funding partners, including 4D Pharma, Abbvie, AcureX, Allergan, Amathus Therapeutics, Aligning Science Across Parkinson’s, AskBio, Avid Radiopharmaceuticals, BIAL, BioArctic, Biogen, Biohaven, BioLegend, BlueRock Therapeutics, Bristol-Myers Squibb, Calico Labs, Capsida Biotherapeutics, Celgene, Cerevel Therapeutics, Coave Therapeutics, DaCapo Brainscience, Denali, Edmond J. Safra Foundation, Eli Lilly, Gain Therapeutics, GE HealthCare, Genentech, GSK, Golub Capital, Handl Therapeutics, Insitro, Jazz Pharmaceuticals, Johnson & Johnson Innovative Medicine, Lundbeck, Merck, Meso Scale Discovery, Mission Therapeutics, Neurocrine Biosciences, Neuron23, Neuropore, Pfizer, Piramal, Prevail Therapeutics, Roche, Sanofi, Servier, Sun Pharma Advanced Research Company, Takeda, Teva, UCB, Vanqua Bio, Verily, Voyager Therapeutics, the Weston Family Foundation and Yumanity Therapeutics. Other data sources we are grateful to have used in this article include the University of Pennsylvania (UPenn) and C-BIG from McGill University (https://zenodo.org/records/17450887).

