## Supplementary Tables and Figures for "Comparison of Automated White Matter Lesion Segmentation Approaches for Use in Large, Multi-Site Data Analyses in Parkinson’s Disease"

**Supplementary Table 1: Performance metrics across the automated approaches in the PD group with medium WML load.**

Values in the upper part of the table are p-values of analysis of variance. The lower part of the table indicates intraclass correlation coefficients and their p-values. Numbers in parentheses indicate the threshold used in the automated algorithms. The asterisk indicates the best value.

| | BIANCA -LOO (0.9) | BIANCA -LOO (0.99) | BIANCA -training (0.9) | BIANCA -training (0.99) | FreeSurfer | FreeSurfer -FAST | LPA (0.4) | LPA (0.5) | LPA -FAST (0.4) | LPA -FAST (0.5) | U-Net | p-value |
|---|---|---|---|---|---|---|---|---|---|---|---|---|
| Dice | 0.58±0.11 | 0.40±0.13 | 0.55±0.12 | 0.39±0.13 | 0.34±0.14 | 0.34±0.15 | 0.41±0.24 | 0.40±0.25 | 0.44±0.21 | 0.41±0.21 | **0.61±0.15*** | < 0.001 |
| Hausdorff | 12.75 ±4.00 | 13.47 ±4.23 | 13.00 ±4.12 | 13.64 ±4.46 | 13.75 ±4.56 | 13.71 ±4.70 | 13.67 ±3.45 | 13.32 ±3.52 | 13.10 ±3.88 | 12.99 ±4.00 | **12.31 ±3.97*** | 0.893 |
| Recall | 0.69±0.19 | 0.53±0.19 | 0.69±0.20 | 0.52±0.20 | 0.22±0.10 | 0.22±0.11 | 0.37±0.29 | 0.33±0.28 | 0.44±0.24 | 0.38±0.23 | **0.76±0.21*** | < 0.001 |
| Precision | 0.34±0.20 | 0.45±0.23 | 0.21±0.15 | 0.32±0.20 | 0.14±0.08 | 0.13±0.09 | 0.30±0.22 | 0.33±0.22 | 0.40±0.19 | 0.44±0.19 | **0.61±0.19*** | < 0.001 |
| F1 score | 0.41±0.19 | 0.43±0.16 | 0.29±0.16 | 0.36±0.16 | 0.15±0.07 | 0.14±0.08 | 0.28±0.18 | 0.26±0.17 | 0.36±0.13 | 0.34±0.13 | **0.64±0.16*** | < 0.001 |
| LOGAVD | 0.34±0.28 | 0.92±0.37 | 0.29±0.26 | 0.86±0.39 | 0.59±0.34 | 0.59±0.34 | 0.42±0.28 | 0.43±0.32 | 0.42±0.28 | 0.48±0.36 | **0.25±0.42*** | < 0.001 |
| *ICC (A, 1)* | | | | | | | | | | | | |
| Coefficient (95% CI) | 0.490 (0.082 -0.731) | 0.114 (-0.052 -0.382) | 0.503 (0.215 -0.707) | 0.134 (-0.061 -0.423) | 0.088 (-0.223 -0.386) | 0.116 (-0.205 -0.414) | 0.335 (0.036 -0.582) | 0.402 (0.103 -0.634) | 0.437 (0.145 -0.659) | 0.434 (0.136 -0.658) | **0.583* (0.310 -0.762)** | |
| p-value | < 0.001 | < 0.001 | < 0.001 | < 0.001 | 0.292 | 0.239 | 0.015 | 0.005 | 0.003 | 0.003 | **< 0.001** | |

**Supplementary Table 2: Performance metrics across the automated approaches in the PD group with high WML loads.**

Values in the upper part of the table are p-values of analysis of variance. The lower part of the table indicates intraclass correlation coefficients and their p-values. Numbers in parentheses indicate the thresholds used in the automated algorithms. The asterisk indicates the best value.

| | BIANCA -LOO (0.9) | BIANCA -LOO (0.99) | BIANCA -training (0.9) | BIANCA -training (0.99) | FreeSurfer | FreeSurfer -FAST | LPA (0.4) | LPA (0.5) | LPA -FAST (0.4) | LPA -FAST (0.5) | U-Net | p-value |
|---|---|---|---|---|---|---|---|---|---|---|---|---|
| Dice | 0.68±0.10 | 0.48±0.12 | 0.66±0.12 | 0.47±0.13 | 0.44±0.11 | 0.46±0.10 | 0.44±0.36 | 0.43±0.35 | 0.37±0.30 | 0.34±0.28 | **0.77±0.07*** | < 0.001 |
| Hausdorff | 16.11 ±7.63 | 19.27 ±8.94 | 16.69 ±8.17 | 19.38± 9.43 | 19.07 ±7.57 | 18.68 ±7.56 | 17.51 ±4.94 | 17.65 ±5.36 | 19.31 ±7.10 | 19.97 ±7.76 | **14.00 ±4.58*** | 0.428 |
| Recall | 0.64±0.16 | 0.45±0.13 | 0.64±0.13 | 0.44±0.14 | 0.20±0.10 | 0.19±0.10 | 0.27±0.18 | 0.23±0.17 | 0.25±0.19 | 0.22±0.17 | **0.85±0.10*** | < 0.001 |
| Precision | 0.44±0.15 | 0.51±0.14 | 0.30±0.14 | 0.38±0.12 | 0.16±0.07 | 0.16±0.08 | 0.28±0.21 | 0.31±0.23 | 0.32±0.24 | 0.38±0.25 | **0.64±0.13*** | < 0.001 |
| F1 score | 0.50±0.13 | 0.47±0.12 | 0.39±0.12 | 0.40±0.11 | 0.17±0.06 | 0.16±0.07 | 0.26±0.17 | 0.24±0.18 | 0.26±0.17 | 0.25±0.17 | **0.72±0.10*** | < 0.001 |
| LOGAVD | 0.40±0.25 | 1.00±0.32 | 0.39±0.28 | 0.99±0.38 | 0.78±0.39 | 0.77±0.30 | 0.23±0.18 | 0.26±0.23 | 0.38±0.29 | 0.52±0.37 | **0.14±0.09*** | < 0.001 |
| *ICC (A, 1)* | | | | | | | | | | | | |
| Coefficient (95% CI) | 0.611 (-0.051-0.870) | 0.234 (-0.103-0.610) | 0.629 (-0.035-0.877) | 0.222 (-0.105-0.594) | 0.302 (-0.114-0.677) | 0.296 (-0.113-0.672) | 0.790 (0.512-0.919) | 0.748 (0.414-0.902) | 0.484 (0.021-0.777) | 0.352 (0.093-0.700) | **0.941* (0.763-0.981)** | |
| p-value | < 0.001 | 0.013 | < 0.001 | 0.018 | 0.008 | 0.008 | < 0.001 | < 0.001 | 0.006 | 0.016 | **< 0.001** | |

**Supplementary Table 3: Dice score and ICC of automated white matter lesion measuring approaches according to lobes in the control group.**

Values are expressed as mean ± standard deviation or number (percentage). The dagger indicates statistical significance. The asterisk indicates the highest value.

| | | BIANCA -LOO (0.9) | BIANCA - LOO (0.99) | FreeSurfer | FreeSurfer -FAST | LPA (0.4) | LPA (0.5) | LPA -FAST (0.4) | LPA -FAST (0.5) | U-Net |
|---|---|---|---|---|---|---|---|---|---|---|
| Frontal | Dice, R | 0.21 ± 0.22 | 0.09 ± 0.12 | 0.05 ± 0.10 | 0.05 ± 0.10 | 0.12 ± 0.19 | 0.1 ± 0.17 | 0.12 ± 0.19 | 0.08 ± 0.15 | **0.34 ± 0.28*** |
| | Dice, L | 0.18 ± 0.20 | 0.08 ± 0.12 | 0.05 ± 0.11 | 0.06 ± 0.12 | 0.09 ± 0.17 | 0.08 ± 0.16 | 0.09 ± 0.15 | 0.06 ± 0.11 | **0.30 ± 0.28*** |
| | *ICC (A, 1)* | 0.526† | 0.231† | 0.287† | 0.202 | 0.394† | 0.309† | 0.129 | 0.089 | **0.881†*** |
| Temporal | Dice, R | 0.01 ± 0.03 | 0.00 ± 0.01 | 0.01 ± 0.04 | 0.01 ± 0.04 | 0.01 ± 0.06 | 0.01 ± 0.04 | 0.01 ± 0.06 | 0.01 ± 0.03 | **0.06 ± 0.17*** |
| | Dice, L | 0.03 ± 0.11 | 0.01 ± 0.04 | 0.02 ± 0.08 | 0.02 ± 0.09 | 0.05 ± 0.15 | 0.04 ± 0.15 | 0.04 ± 0.15 | 0.04 ± 0.14 | **0.07 ± 0.18*** |
| | *ICC (A, 1)* | 0.236† | 0.046 | 0.013 | -0.007 | -0.001 | -0,001 | 0.054 | 0.065 | **0.424†*** |
| Parietal | Dice, R | 0.08 ± 0.15 | 0.03 ± 0.08 | 0.06 ± 0.14 | 0.05 ± 0.13 | 0.09 ± 0.16 | 0.09 ± 0.19 | 0.09 ± 0.17 | 0.08 ± 0.19 | **0.23 ± 0.25*** |
| | Dice, L | 0.15 ± 0.22 | 0.08 ± 0.15 | 0.07 ± 0.17 | 0.06 ± 0.14 | 0.09 ± 0.20 | 0.08 ± 0.19 | 0.09 ± 0.19 | 0.08 ± 0.18 | **0.22 ± 0.28*** |
| | *ICC (A, 1)* | 0.562† | 0.245† | 0.470† | 0.452† | 0.070 | 0.059 | 0.295† | 0.290† | **0.908†*** |
| Occipital | Dice, R | 0.04 ± 0.10 | 0.01 ± 0.04 | 0.02 ± 0.06 | 0.02 ± 0.06 | 0.12 ± 0.21 | 0.11 ± 0.20 | 0.14 ± 0.22 | 0.13 ± 0.21 | **0.13 ± 0.19*** |
| | Dice, L | 0.08 ± 0.17 | 0.01 ± 0.03 | 0.03 ± 0.08 | 0.03 ± 0.08 | 0.09 ± 0.19 | 0.09 ± 0.19 | 0.10 ± 0.21 | 0.11 ± 0.21 | **0.12 ± 0.20*** |
| | *ICC (A, 1)* | 0.374† | 0.029 | -0.003 | -0.018 | 0.062 | 0.067 | 0.271† | **0.385†*** | 0.311† |
| Cingulate | Dice, R | 0.09 ± 0.2 | 0.06 ± 0.14 | 0.04 ± 0.12 | 0.03 ± 0.12 | 0.08 ± 0.19 | 0.07 ± 0.17 | 0.06 ± 0.14 | 0.04 ± 0.12 | **0.10 ± 0.22*** |
| | Dice, L | 0.07 ± 0.17 | 0.02 ± 0.09 | 0.03 ± 0.14 | 0.04 ± 0.14 | 0.05 ± 0.16 | 0.05 ± 0.15 | 0.03 ± 0.09 | 0.02 ± 0.08 | **0.11 ± 0.23*** |
| | *ICC (A, 1)* | 0.562† | 0.245† | 0.470† | 0.452† | 0.070 | 0.059 | 0.295† | 0.290† | **0.908†*** |
| Insular | Dice, R | 0.07 ± 0.18 | 0.04 ± 0.12 | 0.02 ± 0.08 | 0.02 ± 0.1 | 0.05 ± 0.17 | 0.03 ± 0.13 | 0.03 ± 0.14 | 0.02 ± 0.09 | **0.07 ± 0.19*** |
| | Dice, L | 0.09 ± 0.22 | 0.05 ± 0.14 | 0.03 ± 0.12 | 0.03 ± 0.11 | 0.04 ± 0.12 | 0.03 ± 0.11 | 0.03 ± 0.11 | 0.02 ± 0.06 | **0.13 ± 0.26*** |
| | *ICC (A, 1)* | 0.716† | 0.431† | 0.121 | 0.072 | 0.015 | 0.009 | 0.306† | 0.212† | **0.840†*** |
| DWM | Dice, R | 0.44 ± 0.21 | 0.31 ± 0.19 | 0.23 ± 0.20 | 0.24 ± 0.21 | 0.34 ± 0.26 | 0.32 ± 0.26 | 0.35 ± 0.24 | 0.33 ± 0.24 | **0.44 ± 0.25*** |
| | Dice, L | 0.36 ± 0.22 | 0.24 ± 0.17 | 0.20 ± 0.20 | 0.20 ± 0.21 | 0.28 ± 0.25 | 0.26 ± 0.25 | 0.30 ± 0.23 | 0.28 ± 0.23 | **0.41 ± 0.25*** |
| | *ICC (A,1)* | 0.815† | 0.480† | 0.846† | 0.770† | 0.757† | 0.759† | 0.607† | 0.524† | **0.948†*** |

**Supplementary Table 4: Validation of white matter lesion volumes measured by automated approaches using age of participants in the PD group.**
Values are the results of bivariate correlation analysis. The asterisk indicates the highest value.

| | Pearson correlation coefficient | p-value |
|---|---|---|
| Manual segmentation | 0.434 | < 0.001 |
| BIANCA-LOO (0.9) | 0.477 | < 0.001 |
| BIANCA-LOO (0.99) | 0.442 | < 0.001 |
| BIANCA-training (0.9) | 0.486 | < 0.001 |
| BIANCA-training (0.99) | 0.437 | < 0.001 |
| FreeSurfer | 0.409 | < 0.001 |
| FreeSurfer-FAST | 0.371 | < 0.001 |
| LPA (0.4) | 0.487 | < 0.001 |
| LPA (0.5) | **0.489*** | < 0.001 |
| LPA-FAST (0.4) | 0.447 | < 0.001 |
| LPA-FAST (0.5) | 0.445 | < 0.001 |
| U-Net | 0.422 | < 0.001 |

**Supplementary Table 5: Validation of WML volumes measured by automated approaches using Fazekas and Wahlund scores in the PD group.**

Values are the results of partial correlation analysis after adjusting for age. The asterisk indicates the highest value.

| | Fazekas | | Wahlund | |
|---|---|---|---|---|
| | Partial correlation coefficient | p-value | Partial correlation coefficient | p-value |
| Manual segmentation | **0.863*** | < 0.001 | **0.874*** | < 0.001 |
| BIANCA-LOO (0.9) | 0.767 | < 0.001 | 0.766 | < 0.001 |
| BIANCA-LOO (0.99) | 0.761 | < 0.001 | 0.771 | < 0.001 |
| BIANCA-training (0.9) | 0.723 | < 0.001 | 0.737 | < 0.001 |
| BIANCA-training (0.99) | 0.728 | < 0.001 | 0.732 | < 0.001 |
| FreeSurfer | 0.484 | < 0.001 | 0.502 | < 0.001 |
| FreeSurfer-FAST | 0.485 | < 0.001 | 0.495 | < 0.001 |
| LPA (0.4) | 0.729 | < 0.001 | 0.704 | < 0.001 |
| LPA (0.5) | 0.745 | < 0.001 | 0.728 | < 0.001 |
| LPA-FAST (0.4) | 0.703 | < 0.001 | 0.675 | < 0.001 |
| LPA-FAST (0.5) | 0.720 | < 0.001 | 0.697 | < 0.001 |
| U-Net | 0.805 | < 0.001 | 0.811 | < 0.001 |

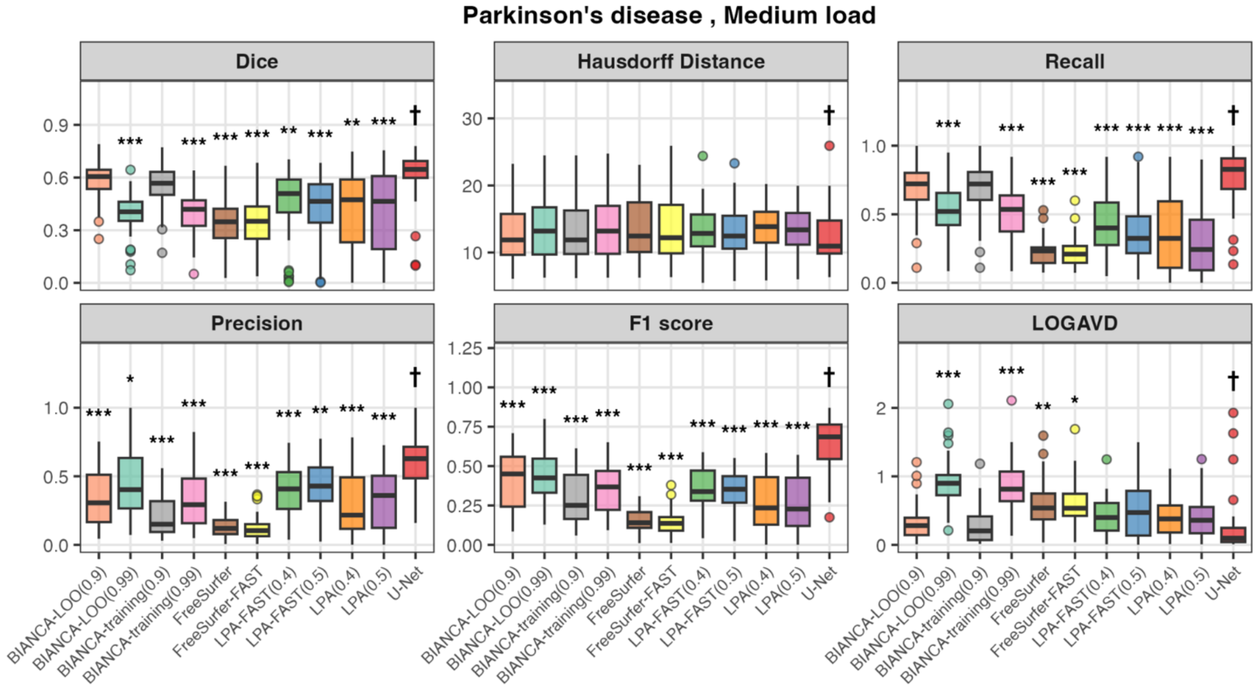


**Supplementary Figure 1: Boxplots of evaluation metrics across the automated approaches in the Parkinson's disease group for medium white matter lesion load.**

Comparison of white matter lesion segmentation algorithm performance in Parkinson's disease patients with medium lesion load (5–15 $cm^3$). Performance of 11 automated segmentation algorithms was evaluated against manual segmentation using six agreement metrics: Dice Similarity Coefficient (higher values indicate better agreement), Hausdorff Distance (lower values indicate better agreement), recall (higher values indicate better agreement), precision (higher values indicate better agreement), F1 score (higher values indicate better agreement), and Log-transformed Absolute Volume Difference (LOGAVD; lower values indicate better agreement). The daggers indicate the best-performing algorithms within each metric. The asterisks indicate statistically significant differences compared to the best-performer (*$p < 0.05$, **$p < 0.01$, ***$p < 0.001$; Tukey HSD test for homogeneous variances or Games-Howell test for heterogeneous variances following Bartlett's test).

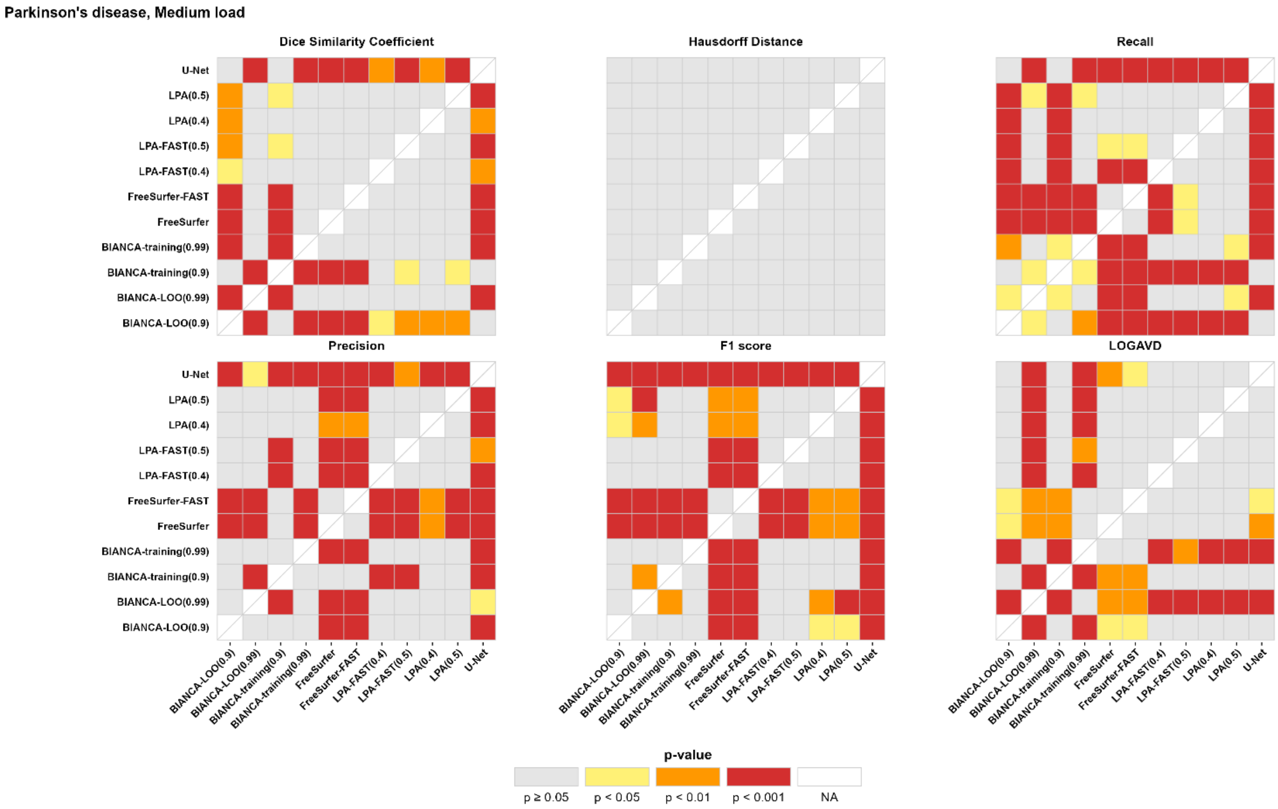


**Supplementary Figure 2: Post-hoc analysis of performance metrics across the automated approaches in the Parkinson's disease group for medium white matter lesion load.**

This figure presents heatmaps of pairwise post-hoc comparisons of automated white matter lesion segmentation algorithms in Parkinson's disease patients with medium lesion load (5–15 cm³). Each cell displays the statistical significance (p-value) of differences in segmentation performance between two algorithms based on six evaluation metrics: Dice similarity coefficient, Hausdorff distance, recall, precision, F1 score, and log-transformed absolute volume difference (LOGAVD). Red indicates greater statistical difference, and white indicates non-significant difference. Post-hoc comparisons were conducted using either Games–Howell or Tukey's honestly significant difference test, depending on the equality of variance.

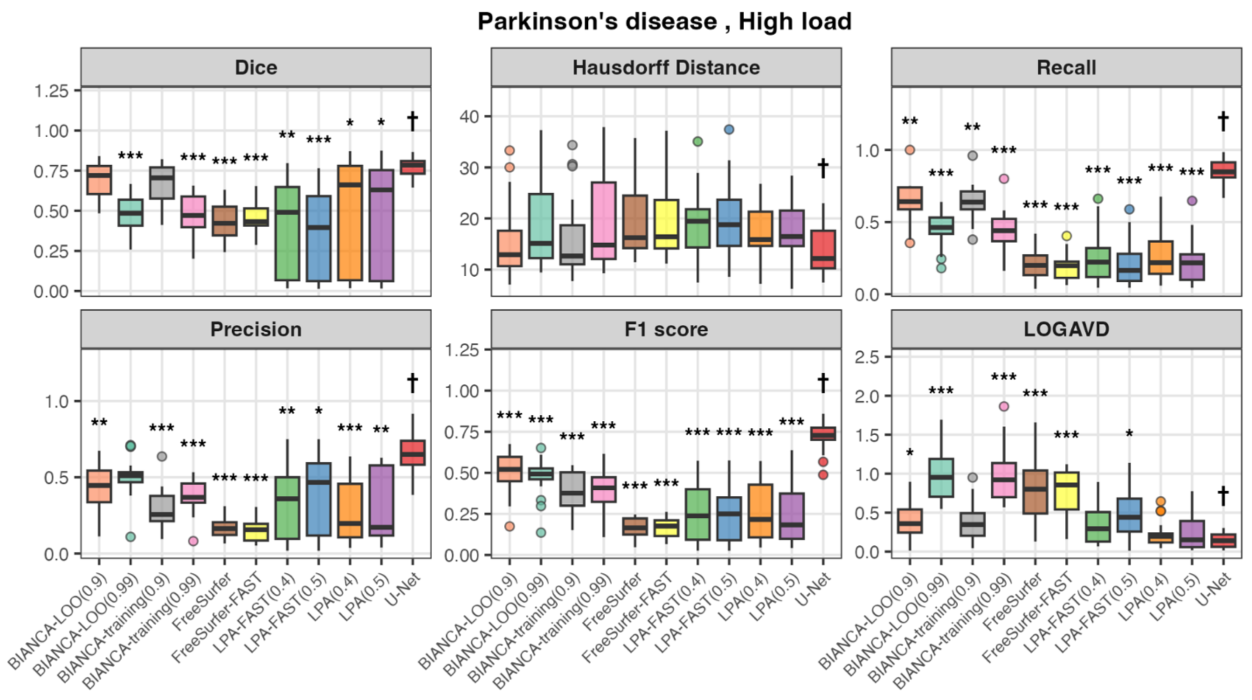


**Supplementary Figure 3: Boxplots of evaluation metrics across the automated approaches in the Parkinson's disease group for high white matter lesion load.**

Comparison of white matter lesion segmentation algorithm performance in Parkinson's disease patients with high lesion load (>15 $cm^3$). Performance of 11 automated segmentation algorithms was evaluated against manual segmentation using six agreement metrics: Dice Similarity Coefficient (higher values indicate better agreement), Hausdorff Distance (lower values indicate better agreement), recall (higher values indicate better agreement), precision (higher values indicate better agreement), F1 score (higher values indicate better agreement), and Log-transformed Absolute Volume Difference (LOGAVD; lower values indicate better agreement). The daggers indicate the best-performing algorithms within each metric. The asterisks indicate statistically significant differences compared to the best-performer (*$p < 0.05$, **$p < 0.01$, ***$p < 0.001$; Tukey HSD test for homogeneous variances or Games-Howell test for heterogeneous variances following Bartlett's test).

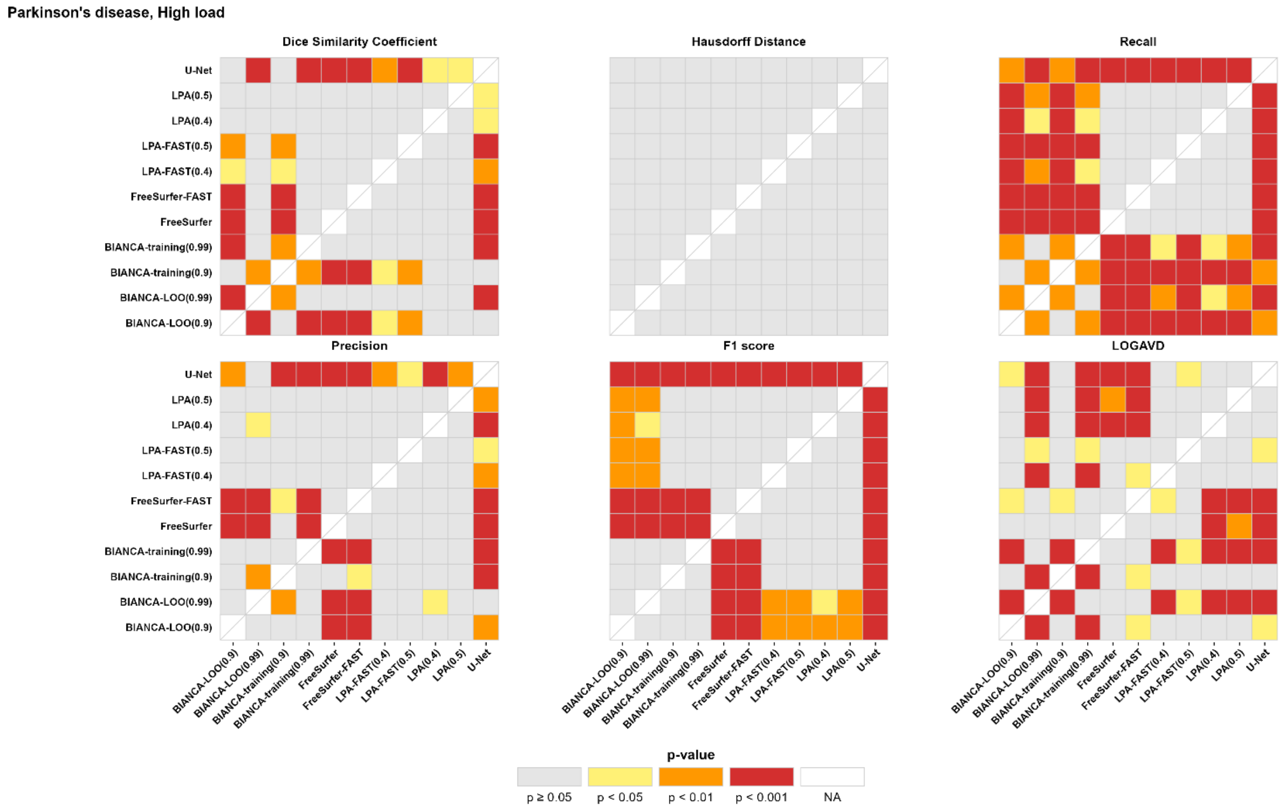


**Supplementary Figure 4:** **Post-hoc analysis of performance metrics across the automated approaches in the Parkinson's disease group for high white matter lesion load.**

This figure presents heatmaps of pairwise post-hoc comparisons of automated white matter lesion segmentation algorithms in Parkinson's disease patients with high lesion load (>15 cm³). Each cell displays the statistical significance (p-value) of differences in segmentation performance between two algorithms based on six evaluation metrics: Dice similarity coefficient, Hausdorff distance, recall, precision, F1 score, and log-transformed absolute volume difference (LOGAVD). Red indicates greater statistical difference, and white indicates non-significant difference. Post-hoc comparisons were conducted using either Games–Howell or Tukey's honestly significant difference test, depending on the equality of variance.
